# Laser Speckle Particle Sizer (SPARSE) Informs the Size Distribution of Tissue Granularities

**DOI:** 10.1101/2024.09.26.615148

**Authors:** Zeinab Hajjarian, Ziqian Zeng, Nichaluk Leartprapun, Seemantini K. Nadkarni

**Affiliations:** Wellman Center for Photomedicine, Massachusetts General Hospital, Harvard Medical School, Boston, MA 02114 USA; Department of Biomedical Engineering, University of Massachusetts Lowell, Lowell, MA 01854 USA

## Abstract

Particle sizing of biomaterials and tissues holds profound implications across diverse industrial and clinical disciplines. Conventional approaches such as dynamic light scattering (DLS) are restricted to dilute liquids, and unable to assess particle size distribution in intact, untampered biomaterials and tissues in their native state. Here, we introduce the laser Speckle PARticle SizEr (SPARSE), which leverages the size-dependent attributes of polarized back-scattered laser speckle to characterize the average radius of a continuum of endogenous scattering particles, in the 10nm-10μm range, in biological fluids and soft tissues in their native state. The SPARSE approach exploits cross- and co-polarized attributes of spatial and temporal laser speckle intensity fluctuations to quantify the average particle size in various specimens, including polystyrene microspheres, milk, whole blood specimens, and intact soft tissues, without prior knowledge or explicit account of refractive index and scattering concentrations. Through scanning the laser beam across the tissue surface, we establish the capability of SPARSE to evaluate a spatial map of heterogenous scattering particle sizes with a resolution of a few 10s of *μ*m that closely mirrors histopathological microstructures in benign and carcinoma breast tissue specimens. By enabling the size characterization of intact tissues and biomaterials across a continuum of granularities, non-destructively in native specimens, SPARSE opens substantial opportunities for quality control in industrial applications, drug development, and advanced clinical diagnostics.

## Introduction

Particle size characterization is an important quantitative target across multiple scientific disciplines, including pharmaceutical research, food processing, and diagnostic pathology^1-5^. In pharmaceuticals, sizing nano drug carriers like liposomes and micelles, a crucial aspect of formulation, involves decomposing the drug bolus into its essential components^2,6^. Similarly, in the food industry, invasive techniques are used for sizing lipid droplets during processes like homogenization and fat reduction, through titrating dairy products in solvents for accurate sizing purposes^1^. Likewise, the diagnosis and management of hematological disorders, including microcyte or macrocyte anemias, thalassemia, sickle cell anemia, myelodysplastic syndromes, and age-related degenerative disorders, hinge on evaluating Mean Corpuscular Volume (MCV) related to the average diameter of red blood cells (RBCs)^5,7-9^. Additionally, grading malignant tumor cells requires visually assessing cell and nucleus size distributions in stained tissue slides, and scrutinizing skewed size distributions of tissue particularities^3^. Therefore, the ability to rapidly assess particle sizes within intact food products, biomaterials, and tissues in their native state offers invaluable opportunities for quality control and diagnostic medicine^10^.

Proximity of optical wavelengths (in the μm range) confers a close relationship between back-scattered light features and particle characteristics, affording the opportunity to reveal the nanostructure and microstructure of minute light-scattering particles^11^. This has led to the emergence of Dynamic Light Scattering (DLS) and Laser Diffraction (LD) as industrial standards for particle sizing. Competent in the sub-micron range, DLS is based on evaluating the intensity fluctuations of back-scattered light, to extract the diffusion coefficient of particle’s Brownian motion, and in turn their size in dilute, low-viscosity liquids^4^. In contrast, LD assesses the angle-dependence of the light diffracted from dispersed particles, and fits it to Mie or Fraunhofer theory for size inference at larger scales, up to a few millimeters^12^. However, the intricate instrumentation and the necessity to conform to the single-scattering approximation via dispersing the particles in low-viscosity solvents bars both techniques from applications in intact biomaterials and tissues. This has created a technological gap in the assessment of particle size within a continuum of granularities in richly scattering specimens in a non-destructive fashion.

To address this gap, we introduce a new technique, the Laser Speckle PARticle SizEr (SPARSE), a non-invasive approach that exploits spatial and temporal intensity fluctuations of back-scattered polarized laser speckle patterns to characterize the size scales of endogenous light-scattering particles in the 10nm-10μm range, within intact biomaterials and soft tissues. When a laser beam is tightly focused on a turbid sample, the back-scattered light exhibits a millimeter-scale intensity envelope that is modulated by the time-varying microscale dark and bright intensity grains, termed speckle^13,14^. Formed because of differences in photon trajectories and optical phases of individual rays, spatio-temporal variations of speckle intensity are exquisitely sensitive to the sample microstructure and dynamics^14-17^. Here we establish that capturing speckle patterns through a linear polarizer, oriented either parallel or perpendicular to the polarization axis of the illumination beam, modifies the temporal fluctuations rate the of speckle grains and the spatial profile of the intensity envelope in concert with the size scale of endogenous light scattering particles. Through polarimetric analysis of the speckle patterns, SPARSE evaluates a set of metrics, integrates them into a simple-to-implement prediction algorithm, and estimates the scattering particle size across a diverse range of biomaterials and tissues. Additionally, by implicitly accounting for variations in optical properties, SPARSE circumvents the need to conform to a single-scattering regimen or prerequisite knowledge of concentration or refractive index variations. We validate the accuracy of SPARSE measurements in biomaterials and biological tissues, exhibiting a wide range of optical scattering and absorption properties, size scales, and polydispersity indices, including polystyrene microspheres, milk phantoms, whole blood, as well as normal and carcinoma-bearing human breast tissue specimens. Additionally, by scanning the focused beam across the sample surface, we demonstrate the capability of SPARSE to map spatially resolved size heterogeneities within both benign and cancerous lesions of the human breast. By facilitating size characterization of intact tissues and biomaterials across a continuous spectrum of particle sizes and optical properties, in a non-destructive fashion, SPARSE presents promising opportunities for diverse applications in both industrial and clinical settings.

## Results

### Principles of SPARSE

Biological tissues consist of nm to μm size particles in constant motion due to the exchange between their kinetic energy and the thermal energy of their microenvironment^15,17^. When a coherent, linearly polarized beam is focused on materials or tissues containing intrinsic light-scattering particles (granularities), photons traverse the illuminated volume along different paths, scatter multiple times, and interfere, forming distinctive laser speckle patterns upon reemerging at the surface^18-21^. These laser speckle patterns exhibit distinct spatial intensity envelopes or shapes, with temporal intensity fluctuations of individual speckle grains (Fig. 1).

**Fig. 1.**
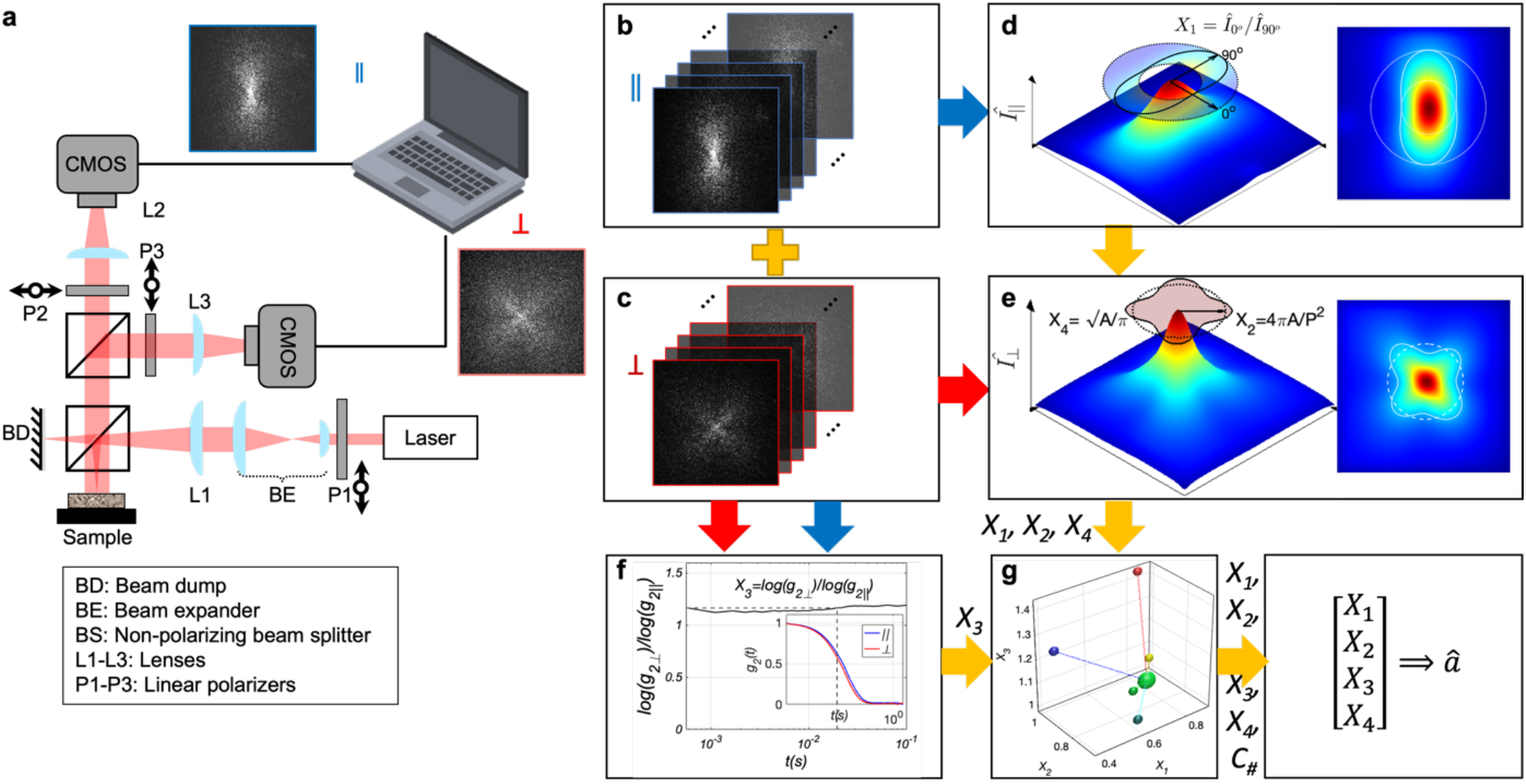
Overview of the SPARSE instrumentation and algorithms. **a** Optical setup of SPARSE. A coherent laser beam is linearly polarized and focused on the luminal surface of the sample. Parallel and perpendicular linearly polarized speckle patterns emanating from the sample are collected by a pair of high-speed CMOS cameras that detect the co-polarized and cross-polarized channels. **b** Co-polarized and **c** Cross-polarized speckle frame series simultaneously collected from a sample. **d** Intensity envelope of the co-polarized speckle frame series, *Î*_∥_. The insets display the top view and the contour of normalized *Î*_∥_ at 30% of the peak intensity. The long axis of the contour is aligned with the polarization axis. To identify the inner and outer circles, the centroid of the contour is determined from the coordinates of the individual points on the contour. The inner and outer circles are then drawn to have their centers at the centroid and pass through the points on the contour that are closest and farthest from the centroid, respectively. Subsequently, *Î*_∥_ is radially averaged in the annular region between the inner and outer circles of the contour, yielding an angular profile, from which 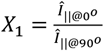 is calculated. **e** Intensity envelope of the cross-polarized speckle frame series, *Î* _*⊥*_. The insets display the top view and the contour of normalized *Î* _⊥_ at 30% of the peak intensity, from which 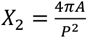 is calculated, where A and P are the area and perimeter of the contour. In addition, *X*_4_=√ (A/π) represents the average radius of the contour. **f** Differential decorrelation rates of co- and cross-polarized speckle time series, defined as 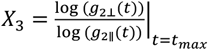 is obtained from the intensity autocorrelation, *g*_*2*_*(t)*, of the acquired speckle time series (inset) at a time instant where the gap is the widest. As seen later, a Monte-Carlo ray tracing algorithm is used to create a synthetic library of *[X*_*1*_,*X*_*2*_,*X*_*3*_, *X*_*4*_] metrics for numerous possible combinations of particle size and optical properties. Cluster analysis of the synthetic *[X*_*1*_,*X*_*2*_,*X*_*3*_, *X*_*4*_] followed by multiple regression analysis provides the particle size estimation equation for each cluster. **G** To estimate the particle size from the experimentally evaluated [*X*_*1*_,*X*_*2*_,*X*_*3*_, *X*_*4*_] metrics, first the Euclidean distance from the cluster centers is evaluated to identify the cluster associated with the experimentally evaluated [X_1_,X_2_,X_3_, X_4_]. Subsequently, the particle size estimation equation corresponding to this nearest cluster center is used to return the average particle size, *a*, using the *[X*_*1*_,*X*_*2*_,*X*_*3*_, *X*_*4*_] metrics.

If granularities in the specimen are smaller than the optical wavelength, *λ*, most photon trajectories involve only a few wide-angle scattering events that largely preserve the incident polarization, forming the parallel or co-polarized component of the speckle pattern^22^. The time-averaged intensity envelope of co-polarized rays, denoted by *Î*_∥_, exhibits a double-lobed profile aligned with the polarization axis^21^. A fainter, more dynamic, cross-polarized speckle is contributed by the longer (and less common) paths that traverse a larger number of scattering events. Therefore, temporal intensity autocorrelation of cross-polarized speckle grains, g_2⊥_(t) decays much faster than co-polarized channel, g_2∥_(t), when particle size scales are smaller than the optical wavelength ^23^.

Conversely, when particle size, *a*, approaches or exceeds *λ*, anisotropic forward-scattering contributes to longer photon trajectories with an equal affinity for co- and cross-polarizations^22^. Consequently, the co-polarized intensity envelope, *Î*_∥_, transforms into a four-leaved pattern, growing a second pair of lobes perpendicular to the polarization axis^21^ (Fig. 1). As such, due to equal affinities to the cross-polarized and co-polarized channels, differences in the intensity autocorrelation decay rates of g_2⊥_ (t) and g_2∥_ (t) are diminished^24^. In contrast to *Î*_∥_, that exhibits a shape change from a dipole to four-leaved, when *a* matches and exceed *λ*, the cross-polarized intensity envelope, *Î*_⊥_ maintains a four-leaved shape, irrespective of *a/λ*. However, the ratio of *Î*_⊥_ area to its perimeter, termed circularity, is reduced. In other words, *Î*_⊥_ evolves from a round to more stellar form, with increasingly distinguished lobes when *a*∼*λ*.

Aside from particle size, the optical properties of tissue also affect both the spatial intensity envelope and temporal dynamics of speckle patterns. Increased particle concentration and refractive index mismatch, *n*_*rel*_*=n*_*particle*_*/n*_*medium*_, foster rich scattering, as quantified by the optical reduced scattering coefficient, *μ*_*s*_*’*^*25*^. Since the mean free path is defined as *l*^*\**^=1/*μ*_*s*_*’*, increased *μ*_*s*_*’* geometrically scales the optical paths. Within speckle field of view (FoV), increasing *μ*_*s*_*’* in turn shrinks the radius of *Î*_∥_ and *Î*_⊥_, and brings forth the longer optical paths with contributions from larger scattering events, promoting a four-leaved *Î*_∥_ shape, and increases the difference in g_2⊥_(t) and g_2∥_ (t) decay rates^19,21^. Conversely, increasing the normalized absorption coefficient, *μ*_*a*_/*μ*_*s*_*’*, i.e. the ratio of absorption and reduced scattering coefficients, prunes the longer paths, pushes the *Î*_∥_ to the double-lobed pattern, and reduces the difference in g_2⊥_ (t) and g_2∥_ (t) decay rates^20,21^. Thus, SPARSE leverages the polarimetric analysis of the spatial and temporal attributes of laser speckle to capture these trends via an array of metrics, denoted as [*X*_*1*_, *X*_*2*_, *X*_*3*_, *X*_*4*_]. These metrics are illustrated in Figs. 1 and 2 and defined in detail in the Methods section. Briefly, *X*_*1*_ captures the azimuth angle variation of *Î*_∥_ as 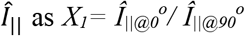. Here 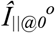 and 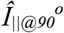 represent the radially average *Î*_∥_ at 0° and 90° azimuth angle, in the annular region between inner and outer circles of its contour at 30% of the maximum intensity. The circularity of *Î*_⊥_ is quantified by 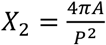, where *A* and *P* are the area and the perimeter of the *Î*_⊥_ contour. In addition, the differential decorrelation rate of g_2⊥_(t) and g_2∥_(t) is captured by 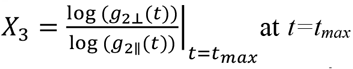 at *t=t*_*max*_, where the slopes of the two curves and in turn their gap was maximized. Finally, the radial extent of *Î*_⊥_is captured by *X*_*4*_*=*√*(A/π)*. These metrics are then incorporated within a prediction algorithm (described below) to estimate the scattering particle size, *a* (Fig. 1). We had previously demonstrated the proof of concept for comparing the azimuth angle profile of *Î*_∥_ with a look up table (LUT) to estimate the size^21^. Based solely on the spatial profile of the co-polarized speckle intensity envelope, this prior work remained limited to semi-quantitative, discrete size estimating over ∼250nm-2.5μm range (single decade), in purely scattering samples of limited turbidity, and overlooked the impact of absorption and refractive index variations. In what follows we detail how SPARSE leverages the full extent the spatio-temporal speckle statistics to provide quantitative evaluating of size, over three decades of scales, 10nm-10μm, in biological samples, irrespective of optical properties.

**Fig. 2.**
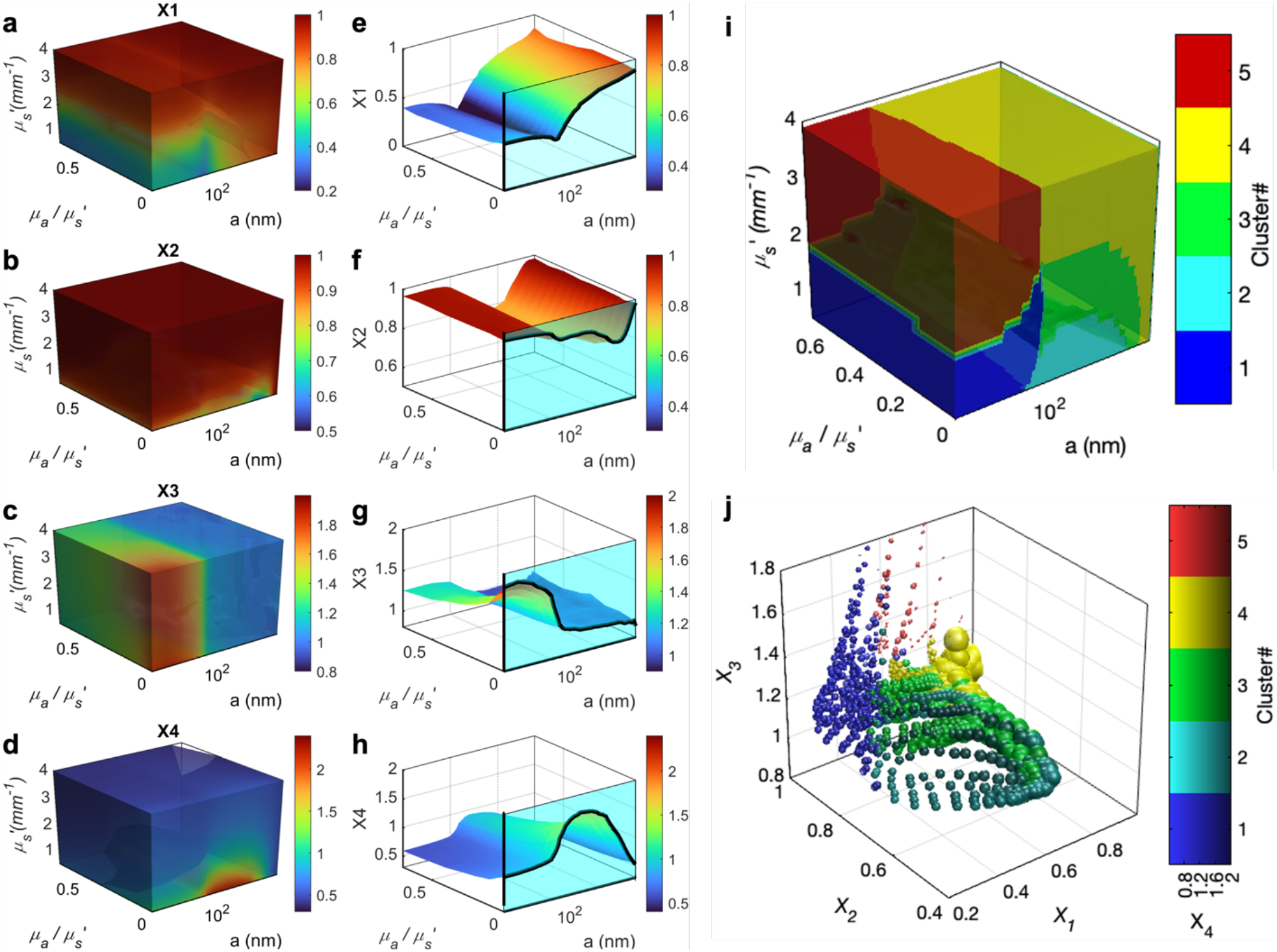
Synthetic library of size-dependent metrics, created by PSCT-MCRT. **a-d** 3D plots defining the associations between each of the metrics *X*_*1*_, *X*_*2*_, *X*_*3*_, *X*_*4*_, *μ*_*s*_*’*, and *μ*_*a*_*/μ*_*s*_*’* for a range of particle radii, *a* (10nm-10μm). **e-h** An example of a single cross-sectional plane of the 3D plots (for *μ*_*s*_*’*=1 mm-1 typical of biological tissue) showing the associations between *X*_*1*_, *X*_*2*_, *X*_*3*_, *X*_*4*_, with particle size and *μ*_*a*_*/μ*_*s*_*’* variations. The line plots, highlighted by a cyan background corresponds to *μ*_*s*_*’*=1 mm^-1^ and *μ*_*a*_=0. It is evident that *X*_*1*_ shows a slight decrease as *a* increases from 10-100 nm, but increases rapidly in the a:125nm-10μm range. On the other hand, X_2_ presents an absolute minimum at the sizes closer to the laser wavelength. *X*_*3*_ presents a decaying trend in *a*, whereas *X*_*4*_ resembles an inverted U shape, almost reverse of *X*_*2*_, with a peak that is slightly shifted towards smaller sizes, compared to the valley in X_2_ curve. The size-dependent variations of *X*_*1*_, *X*_*2*_, *X*_*3*_, *X*_*4*_ are reduced by absorption, as evidenced by diminished peaks and valleys in the opposite wall plots. **i** Clustering analysis of *[X*_*1*_,*X*_*2*_,*X*_*3*_, *X*_*4*_] metrics partitions the [*a, μ*_*a*_, *μ*_*s*_*’*] space to 5 distinct clusters. **j** Scatter plot of the cluster assignments in the *[X*_*1*_, *X*_*2*_, *X*_*3*_, *X*_*4*_] space. The 3 axes of the plot correspond to *X*_*1*_, *X*_*2*_, and *X*_*3*_, while *X*_*4*_ is depicted by varying the luminescence of the cluster color. The sizes of spherical markers are proportionate to the particle volume, i.e. ^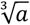^. A tailored regression model that formulates *a* as a function of *X*_*1*_, *X*_*2*_, *X*_*3*_, *X*_*4*_ is obtained for each cluster.

### SPARSE Prediction algorithm

To develop the SPARSE particle size estimation algorithm, we expanded our previous polarization-sensitive Monte-Carlo ray tracing approach, to simulate both the diffused light intensity and its correlation-transfer, to obtain not only the first-(i.e. average spatial intensity) but also the second-order (i.e. temporal intensity auto-correlation) statistics of both co- and cross-polarized speckle. This permitted simulating both the polarized intensity envelopes *Î*_∥_, *Î*_⊥_ and polarized intensity temporal autocorrelation curves g_2∥_(t), and g_2⊥_(t) in turbid media of biologically relevant optical properties *μ*_*s*_*’*: 0.25-4 mm^-1^, and *μ*_*a*_: 0-70%μ_s_’, over scattering size range of *a*=10nm-10*μm* (See Supplementary Materials)^26^. It was assumed that *n*_*rel*_*=1*.*1*, and *μ*_*s*_*’* variations were achieved solely by modifying the particle size and concentration and not the *n*_*rel*_. We then reconstructed the *Î*_∥_, *Î*_⊥_, g_2∥_(t), and g_2⊥_(t) from the simulated photon trajectories as detailed in Supplementary Materials.

From these from simulated polarized speckle attributes, a synthetic library of *[X*_*1*_, *X*_*2*_, *X*_*3*_, *X*_*4*_] metrics was calculated for an expansive set of [*a, μ*_*a*_, *μ*_*s*_*’*], covering a significantly larger size range, in turbid media of arbitrary optical properties as illustrated in Fig. 2. The simulated *[X*_*1*_, *X*_*2*_, *X*_*3*_, *X*_*4*_] metrics served as quantitative predictors of particle size over 3 decades, while also implicitly accounting for turbidity variations, as depicted in Fig. 2. As an example, for *μ*_*s*_*’=1 mm*^*-1*^ *and μ*_*a*_ =0, our simulation results provide a detailed account of *Î*_∥_ variations with size, demonstrating that when *a* ∼10s nm, *Î*_∥_ is almost elliptical with a subtle notch in the midline that deepens near 100nm, causing *X*_*1*_ to decrease with *a* in this range. A second pair of lobes raise above 150 nm, leading to the growth of *X*_*1*_ with *a*, before it saturates beyond 10 μm (Fig. 2e, line plots). Our results also showed that *X*_*2*_ remains close to 1, for sizes much smaller or larger than *λ*, but reduces significantly near *λ* (Fig. 2f, line plot). The precise location of this absolute minimum changes with the scattering asymmetry parameter, *g*, and in turn *a* and *n*_*rel*_. *X*_*3*_ is consistently decreasing with *a*, but exhibits a significantly steeper slope in submicron range (Fig. 2g, line plot). X_4_ resembles an inverted U shape, exhibiting an almost opposite trend relative to X_2_. However, the location at which X_4_ is maximized is slightly shifted towards smaller sizes, compared to the valley in *X*_*2*_ curve. The line plots detailed above correspond to *μ*_*s*_*’=1 mm*^*-1*^ *and μ*_*a*_ =0. Similar line plots may be obtained by referring to cross sections of Fig. 2 (a-d), at the intersection of the desired *μ*_*s*_*’ and μ*_*a*_ combination. These line plots exhibit different increasing or declining trends for *[X*_*1*_, *X*_*2*_, *X*_*3*_, *X*_*4*_] with particle radii, and the specific location and magnitude of the maxima and minima are additionally modulated by *μ*_*a*_ and *μ*_*s*_*’*. For instance, increasing the *μ*_*a*_ */μ*_*s*_*’* tends to diminish the *[X*_*1*_, *X*_*2*_, *X*_*3*_, *X*_*4*_] variations by size, as evidenced by cross-sections parallel to the cyan-filled wall nearing the opposite wall.

From the above trends, it is evident that the mathematical forms relating *[X*_*1*_,*X*_*2*_,*X*_*3*_, *X*_*4*_] metrics to *a* vary both across the size range and in response to optical properties, suggesting that multiple particle size estimation equation may be needed within different regions of [*a, μ*_*a*_, *μ*_*s*_*’*] and in turn *[X*_*1*_, *X*_*2*_, *X*_*3*_, *X*_*4*_] space. Cluster analysis of synthetic *[X*_*1*_, *X*_*2*_, *X*_*3*_, *X*_*4*_] metrics reveal that this space may be divided to an optimum number of 5 clusters. Subsequently, a step-wise regression analysis of *[X*_*1*_, *X*_*2*_, *X*_*3*_, *X*_*4*_] within individual clusters yields a tailored particle size estimation equation expressing *a* as a function of *[X*_*1*_, *X*_*2*_, *X*_*3*_, *X*_*4*_], belonging to that cluster (Supplementary methods, eqns. 10-14). Therefore, to estimate the particle size from experimentally measured polarized speckle frame series, we simply need to follow the steps outlined in the flowchart of Fig. 1, to calculate the experimental *[X*_*1*_, *X*_*2*_, *X*_*3*_, *X*_*4*_] metrics, identify the cluster to which the sample belongs, and plug in the metrics in the corresponding particle size estimation equation (Supplementary Materials, eqn. 10-14), and obtain *a*. Details of the clustering analysis and the particle size estimation equations are elaborated in the “Methods” and “Supplementary Materials”. In what follows, we investigate and validate the SPARSE approach by applying it for particle size measurement in monodispersed suspensions and biological tissues of increasing complexities.

### Characterizing the mono-dispersed polystyrene microsphere suspensions

We first evaluated mono-dispersed polystyrene microsphere suspensions in aqueous glycerol solutions to assess the accuracy of the SPARSE approach to measure *a* in the 50 nm to 5 μm range, as illustrated in Figure 3. The refractive indices for the microspheres and glycerol solution are *p*_*article*_=1.59 and *n*_*medium*_≈1.45, respectively, resulting in *n*_*rel*_=1.1, which falls within the biologically relevant range^27^. This is the same *n*_*rel*_ used to develop the synthetically generated *[X*_*1*_, *X*_*2*_, *X*_*3*_, *X*_*4*_] library and the SPARSE prediction algorithm. Figure 3 shows *Î*_∥_, *Î*_⊥_, g_2∥_(t), and g_2⊥_(t) obtained experimentally using the SPARSE instrument (Fig. 1A) for a selected subset of polystyrene microsphere suspensions with *a* of 75 nm, 100 nm, 250 nm, and 5 μm (*μ*_*s*_*’* was 0.77, 1.08, 0.97, and 0.63 mm^− 1^, respectively). As *a* increases, *X*_*1*_ increases from 0.36 to 0.95, while *X*_*2*_ shows less variation at these selected sizes, except for the lower value at 75 nm. *X*_*3*_ and *X*_*4*_ exhibit the expected decreasing and concave trends, respectively (similarly observed in synthetic data Fig. 2). SPARSE holds the promise to estimate *a* regardless of *μ*_*s*_*’* variations. The experimentally determined *[X*_*1*_, *X*_*2*_, *X*_*3*_, *X*_*4*_] metrics of polystyrene bead are compared to the cluster centers to identify the one with the shortest Euclidean distance. Substituting the *[X*_*1*_, *X*_*2*_, *X*_*3*_, *X*_*4*_] metrics in the particle size estimation equation of corresponding clusters yields the predicted *a* value. A significant correlation is observed between SPARSE and DLS measurements (r=0.92, p<10^−4^, intercept=24 nm) across the entire 50nm-5*μ*m polystyrene bead sizes. Bland-Altman analysis indicates mean difference of 80 nm (95% CI: -750-910 nm) between DLS and SPARSE measurement. Also noted in the scatter plot is the deviation of SPARSE from DLS for *a*< 100 nm, which is likely caused by significant dependence of particle size estimation equation on *X*_*3*_ in this size range, and in turn the critical modulation of this metric by both *a* and *μ*_*s*_*’*, at the scale of few 10s of nm, as discussed later. We have summarized SPARSE and DLS measurement results for all the polystyrene microspheres, along with theoretically calculated *μ*_*s*_*’* values, in Supplementary Table S3.

**Fig. 3.**
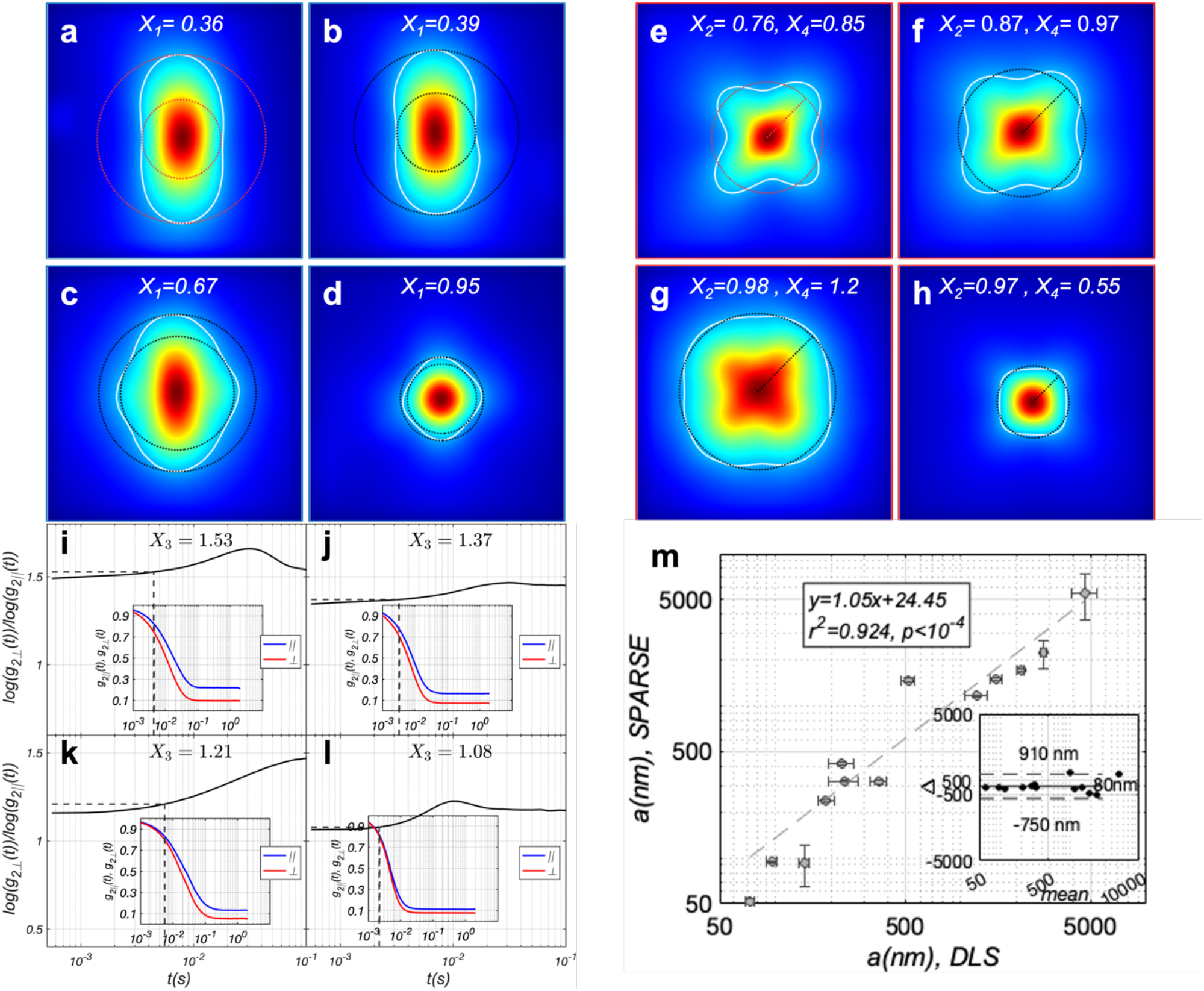
Sizing polystyrene microspheres. **a-d** *Î*_∥_ corresponding to aqueous glycerol suspensions of polystyrene microspheres exhibiting radii of 75nm, 100nm, 250 nm, and 5000 nm, respectively. Also displayed in each panel is the contour of *Î*_∥_ at 30% of its maximum together with the calculated *X*_*1*_ values. **e-g** Corresponding *Î*_⊥_ of polystyrene microsphere phantoms. Similarly, the contour at 30% is traced and the calculated *X*_*2*_ and *X*_*4*_ are displayed. **i-l** Ratio of the speckle decorrelation rate in perpendicular and parallel polarization, defined as log(g_2⊥_(t))/log(g_2∥_(t)). The inset displays g_2⊥_(t) in red and g_2∥_(t) in blue. *X*_*3*_ is calculated at the temporal point where the g_2_(t) slope is maximized (dashed lines in the main plot and inset). **m** Scatter diagram of particle radius, *a*, predicted by SPARSE exhibits a strong, statistically significant correlation with standard DLS measurements. Bland-Altman analysis indicates mean difference of 80 nm (95% CI: -750-910nm) between DLS and SPARSE measurement.

### Characterizing the particle size in milk specimens of varying fat content

Particle sizing of milk is an integral part of quality control in the dairy industry^1^. It has been reported that in homogenized milk, proteins like casein micelles exhibit typical radii of 100 nm, while fat globules span the 200 nm to 2 μm range^28^. We tested the utility of SPARSE for particle sizing in commercially available non-fat (0%), low-fat (1%), reduced-fat (2%), and whole milk (4%), with unknown optical properties and refractive index variations. Our measurements indicate that by increasing fat content from 0% to 4%, *X*_*1*_ increases from 0.44 to 0.59, *X*_*2*_ increases from 0.77 to 0.99, and *X*_*3*_ consistently decreases from 1.45 to 1.01. *X*_*4*_ increases from 0.69 in 0% fat milk to 0.94 in 1% milk, but then reduces to 0.9 and 0.77 in 2% and 4% fat milk, respectively. This is likely due to the competing effects of simultaneous increase in *a* and *μ*_*s*_*’* with fat content, in respectively raising and reducing *X*_*4*_, in this range. (Fig. 4). Based on these metrics, SPARSE estimates average particle sizes of 99.6, 105, 142, and 180 nm for 0%, 1%, 2%, and 4% milk samples, which strongly and significantly correlate with DLS measurements. Bland-Altman analysis suggests a mean difference of 12 nm (95%CI -8.7-33.6nm) between DLS and SPARSE measurement. Deviations are larger for 1% and 2% fat milk, likely because size differences are in the order of few 10s of nm, approaching the resolution limit of SPARSE and DLS. These estimates also align with the expected trend in particle sizes based on the proportions of protein micelles and fat globules^29^. The reported values for the refractive indices of fat globules (*n*_*f*_=1.46, *n*_*f*_*/n*_*w*_=1.1) and protein micelles (*n*_*p*_*=1*.*57, n*_*p*_*/n*_*w*_=1.18) are either comparable or higher than those for polystyrene microsphere phantoms^29,30^. Here, *n*_*w*_=1.33 represents the refractive index of water, which constitutes most milk content. Nevertheless, SPARSE’s prediction algorithm remains applicable to these biologically relevant samples.

**Fig. 4.**
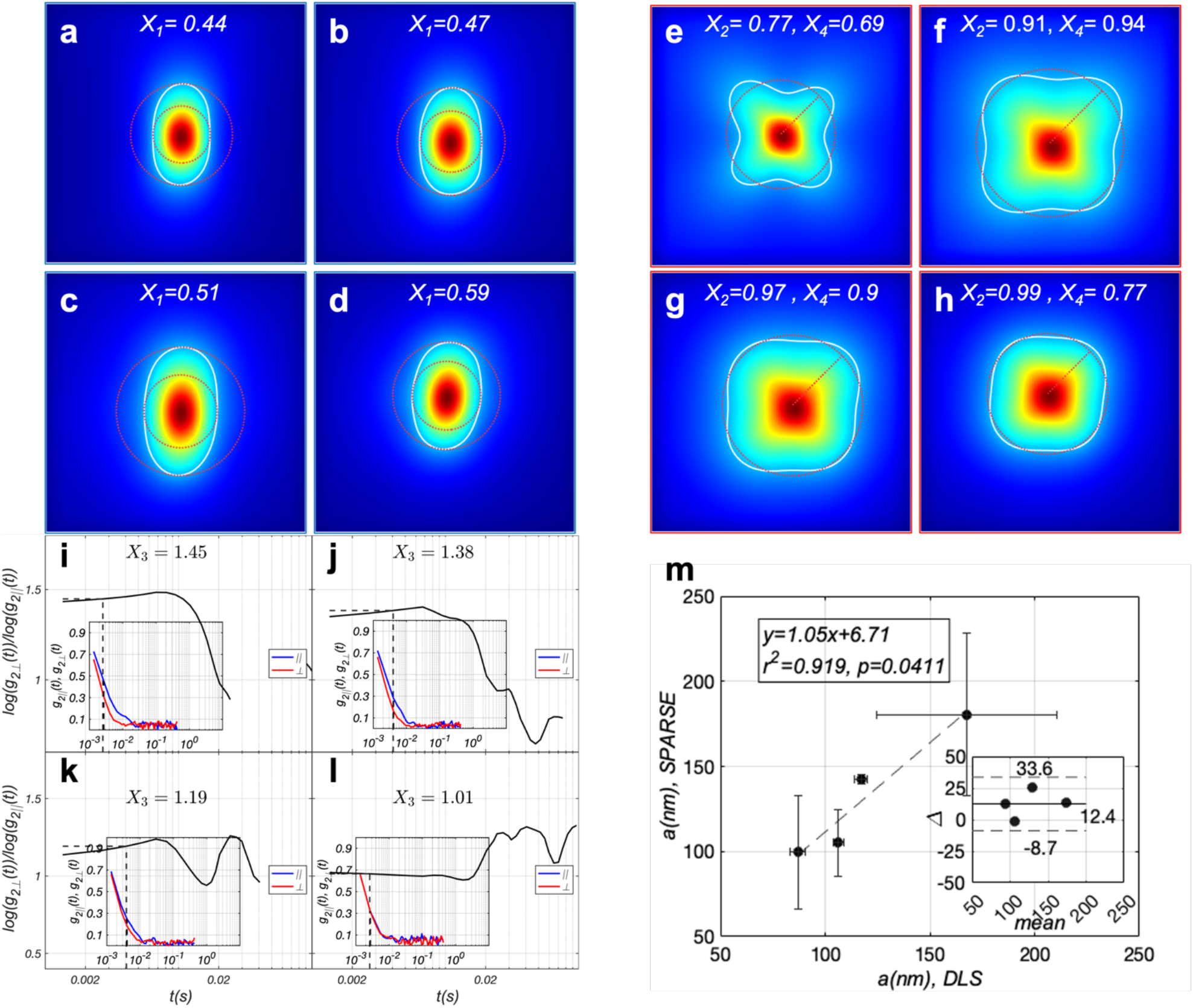
Characterizing the particle size in milk. *Î*_∥_ of **a** non-fat, **b** low-fat (1%), **c** reduced-fat (2%), and **d** whole (4%) milk, respectively. Also displayed in each panel is the contour of *Î*_∥_ at 30% of its maximum together with the calculated *X*_*1*_ values. **e-g** Corresponding *Î*_⊥_ of milk. Similarly, the contour at 30% is traced and the calculated *X*_*2*_ and *X*_*4*_ are displayed. **i-l** Ratio of the speckle decorrelation rate in perpendicular and parallel polarization, defined as log(g_2⊥_(t))/log(g_2∥_(t)). The inset displays g_2⊥_(t)in red and g_2∥_(t) in blue. *X*_*3*_ is calculated at the temporal point where the g_2_(t) slope is maximized, as indicated by a dashed line in both the main plot and the inset. **m** Scatter diagram of *a*, predicted by SPARSE exhibits a strong, statistically significant correlation with standard DLS measurements. Bland-Altman analysis shows a mean difference of 12 nm (95%CI -8.7-33.6nm) between the two measurements.

### Tracing the shrinkage of RBC size in whole blood samples of increased tonicity

Sizing red blood cells (RBCs) is essential for diagnosing a variety of anemias and other blood disorders that result in microstructural changes in RBCs^7^. Here, we tested the capability of SPARSE to quantify RBC size in fresh whole-blood and trace changes in RBC size. Compared to microspheres and milk, which presented negligible absorption, at λ =633 nm blood exhibits not only scattering, but also absorption properties due to its hemoglobin content, with *μ*_*s*_*’*=1.2 *mm*^*-1*^, and *μ*_*a*_*=0*.*3 mm*^*-1*^as reported in the literature

Blood was obtained from a healthy human donor and its tonicity was modified by spiking 50μl of blood with 10μl of saline of 0.9-20% NaCl concentrations, achieving a final NaCl concentrations of 0.9%-4.08%, in blood^32^. Increased osmolarity caused RBC shrinkage, while at the same time changed the conformations of heme groups in RBC, simultaneously reducing *μ*_*a*_ at the illumination wavelength^32,33^. Progressive increase in RBC density also likely enhanced the *n*_*rel*_ and in turn *μ*_*s*_^’34^. This was evidenced by the brighter red hues observed in samples with higher NaCl concentrations (Supplementary Materials), presenting an opportunity to assess the feasibility of applying SPARSE to track changes in particle size with concurrent changes in optical properties in biological samples. Figure 5 depicts *Î*_∥_, *Î*_⊥_, g_2∥_(t), and g_2⊥_(t) for blood samples of 0.9%, 1.42%, 1.75%, and 2.42% final NaCl concentration, together with the traces of estimated *a* versus NaCl concentration.

**Fig. 5.**
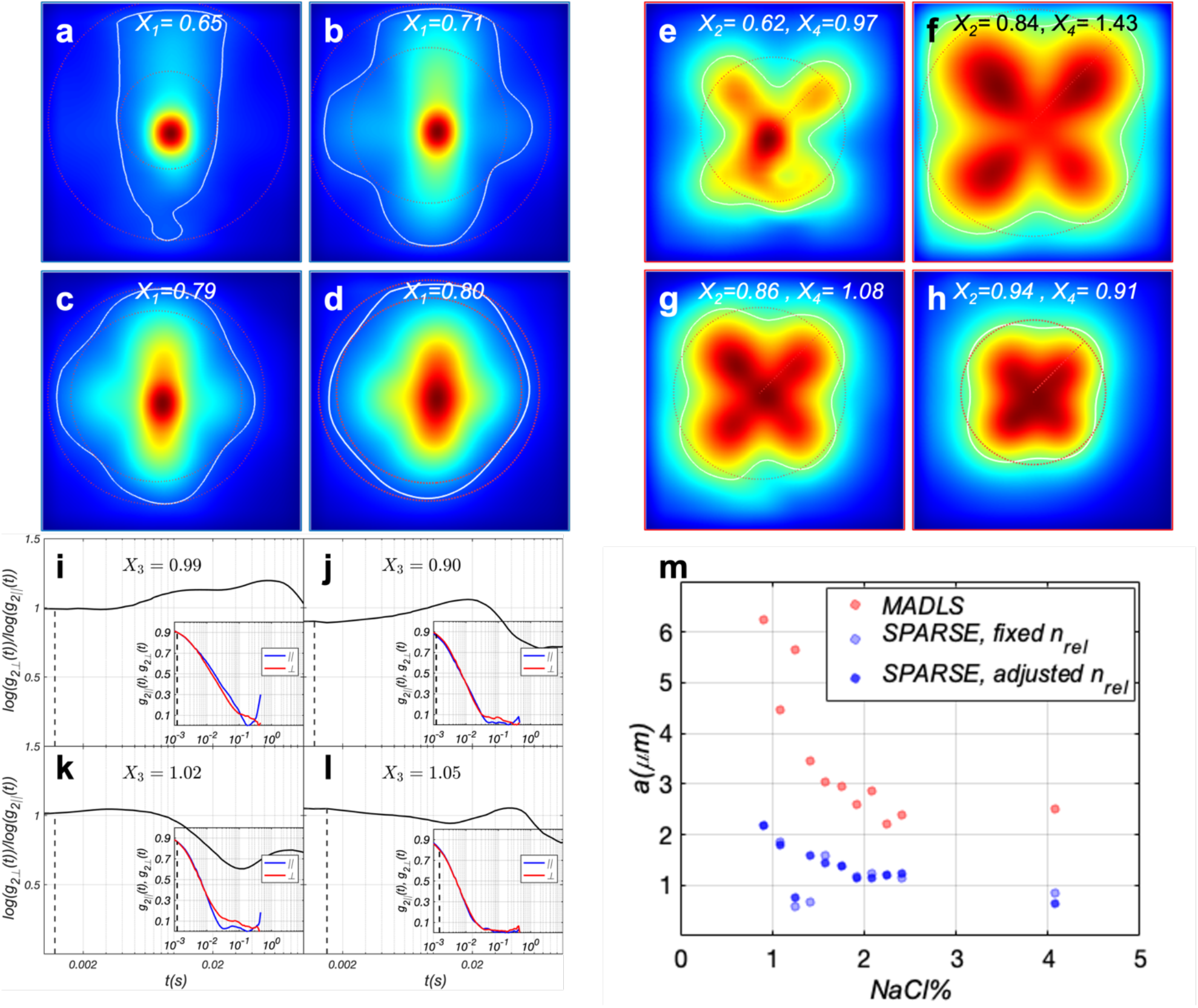
Tracing RBC shrinkage in response to increased saline concentration of plasma. *Î*_∥_ of whole blood samples of **a** 0.9%, **b** 1.42%, **c** 1.75%, and **d** 2.42% NaCl concentration, respectively. Also displayed in each panel is the contour of *Î*_∥_ at 30% of its maximum together with the calculated *X*_*1*_ values. **e-g** Corresponding *Î*_⊥_ of whole blood samples. Similarly, the contour at 30% is traced and the calculated *X*_*2*_ and *X*_*4*_ are displayed. **i-l** Ratio of the speckle decorrelation rate in perpendicular and parallel polarization, defined as log(g_2⊥_(t))/log(g_2∥_(t)). The inset displays g_2⊥_(t) in red and g_2∥_(t) in blue. *X*_*3*_ is calculated at the temporal point where the g_2_(t) slope is maximized, as marked by the dashed line in the main graph and the inset. **m** Particle radius, *a*, predicted by SPARSE, before and after correcting for refractive index variations. Trends agree with the expectations and multi-angle DLS (MADLS) measurements. Given the discoid shape of RBCs and polydisperse nature of the blood, MADLS (vs DLS) was deemed more suitable for benchmarking SPARSE in blood.

The *Î*_∥_ of isotonic blood appeared elliptical, exhibiting X_1_=0.65, which is lower compared to polystyrene phantoms of comparable radii (2.78 μm). Our MCRT simulations suggested that this trend is likely because of significant *μ*_*a*_ of at least *0*.*3 mm*^*-1*^ of blood at 633 nm (Supplementary Materials)^31^. As saline concentration increased, secondary lobes of *Î*_∥_ began to appear and X_1_ raised to 0.8, likely due to reduced *μ*_*a*_ and enhanced *μ*_*s*_*’*, concurrent with reduced RBC size. This is corroborated by *Î*_⊥_, X_2_ and X_4_ trends, which revealed the expansion of speckle intensity envelope, suggesting a reduction in *μ*_*a*_ prior to *μ*_*s*_*’* enhancement. Specifically, a small increase in tonicity expanded the *Î*_⊥_ and made it more circular because longer optical paths that once were absorbed were now emerging at peripheral regions. Also, in contrast to what was observed in polystyrene phantoms and milk, *Î*_⊥_ was maximized in the center of lobes (i.e. leaflets) instead of the envelope centroid. Our MCRT simulations attribute this phenomenon to the lower *n*_*rel*_=1.03 of whole blood (n_RBC_=1.402, n_plasma_=1.36, Supplementary Methods) compared to that used to develop the synthetic library^35^. With higher tonicity, RBCs contracted, causing *Î*_⊥_ to shrink, shifting its maxima toward the centroid, increasing X_2_, and reducing X_4_, in response to increased *n*_*rel*_, and in turn *μ*_*s*_*’*. As expected, *X*_*3*_ remained close to unity, yet both g_2∥_(t), and g_2⊥_(t) decayed faster at higher NaCl concentrations, reflecting a combination of size reduction, reduced absorption, and enhanced scattering^20^. The metrics *[X*_*1*_, *X*_*2*_, *X*_*3*_, *X*_*4*_] were used to identify the most nearby cluster, and were then substituted in the particle size estimation equation corresponding to that cluster to obtain *a*. Our results demonstrate that SPARSE accurately traces RBC shrinkage, in concert with MADLS trends. Given the discoid shape of RBCs and the polydisperse nature of blood, multi-angle DLS (MADLS) measurements were used to benchmark the SPARSE results. SPARSE’s absolute values were in the vicinity of 2.8 μm, reported as the sphere-equivalent radius of RBCs in the literature^36^. In contrast, MADLS reported a larger particle size, likely due to discoid shape of RBCs and the offset of results towards the longer radius in the order of 4μm together with significant polydispersity for blood specimens. Deviations of SPARSE results from the expected trend at 1.25% and 1.42% NaCl, were likely caused by a significant reduction of *n*_*rel*_, at these concentration. To ameliorate the reduced accuracy induced by *n*_*rel*_ variations, an additional synthetic library and the corresponding particle size estimation equations were developed assuming *n*_*rel*_=1.03 (Supplementary Methods). This alternative model was used whenever the radial distance of *Î*_⊥_ maxima from the centroid was over 50% of X_4_. Adjusting for index variations improved the SPARSE results at 1.25% and 1.42% NaCl concentrations (Fig. 5), with little influence at other concentrations. Nevertheless, a full recovery of the trends at 1.25% remained out of reach, due to noisy g_2_(t) curves, which artificially increased X_3_ to above 1, causing the estimated a to drop at this concentration.

### Mapping particle size distributions in heterogenous benign and cancerous tissue specimens

Our results in polystyrene microbeads, milk, and whole blood above demonstrated the capability of SPARSE in evaluating the average particle size in homogenous fluid suspensions that exhibit a range of *n*_*rel*_ and particle concentrations, and in turn optical properties, including with high optical absorption. Next, we tested the capability of the SPARSE approach for particle sizing in heterogenous breast tissue specimens, by scanning the laser beam across the sample and measuring local polarization attributes of speckle patterns to ultimately reveal colormaps of particle radii, *a*, distributions (Figs. 6 and 7).

**Fig. 6.**
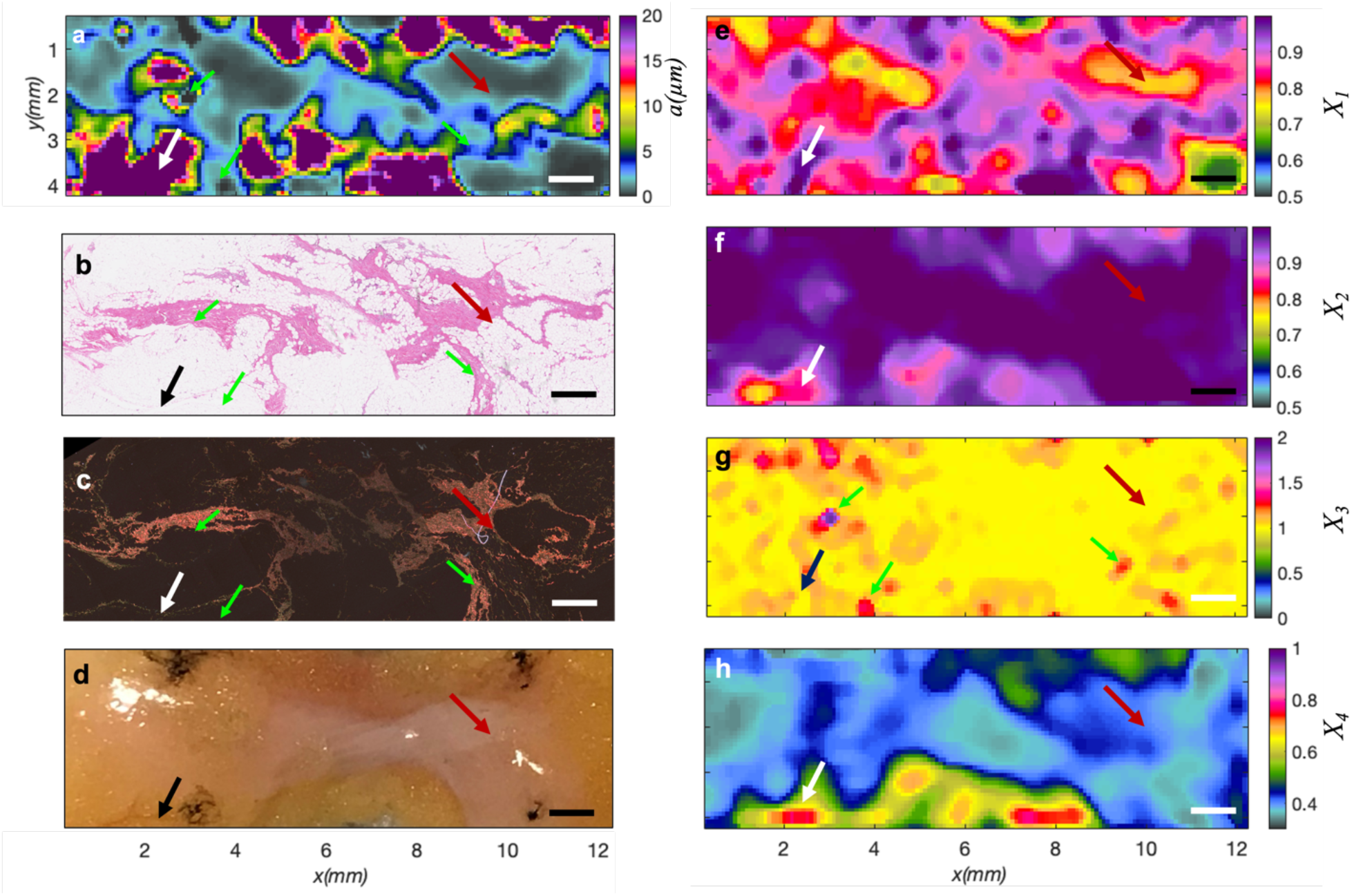
Particle size mapping in benign fibroadipose breast tissue. **a** SPARSE map of *a*. Regions of increased particle size correspond to fat globules in breast tissue (black or white arrows), whereas areas of reduced size align with fibrous structures (maroon arrows). **b** H&E section shows the fibrous (maroon arrow) and adipose (black arrow) regions. **c** PSR section shows corresponding collagen-rich regions. **d** Photograph displaying the gross pathology of fibroadipose tissue. **e** *X*_*1*_ is slightly lower in the regions that correspond with fibrous tissue (maroon arrow) compared to adipose regions (white). **f** *X*_*2*_ on the other hand, exhibits slightly higher values in fibrous areas. **g** Spatial map of *X*_*3*_ is nearly 1 across the tissue except for isolated specks of higher values that occasionally coincide with fibrous structures (green arrows), likely due to significant thinning of collagen fibers in these areas. **h** *X*_*4*_ values are lower in the fibrous regions compared to surrounding adipose tissue. Scale bars are 1 mm.

**Fig. 7.**
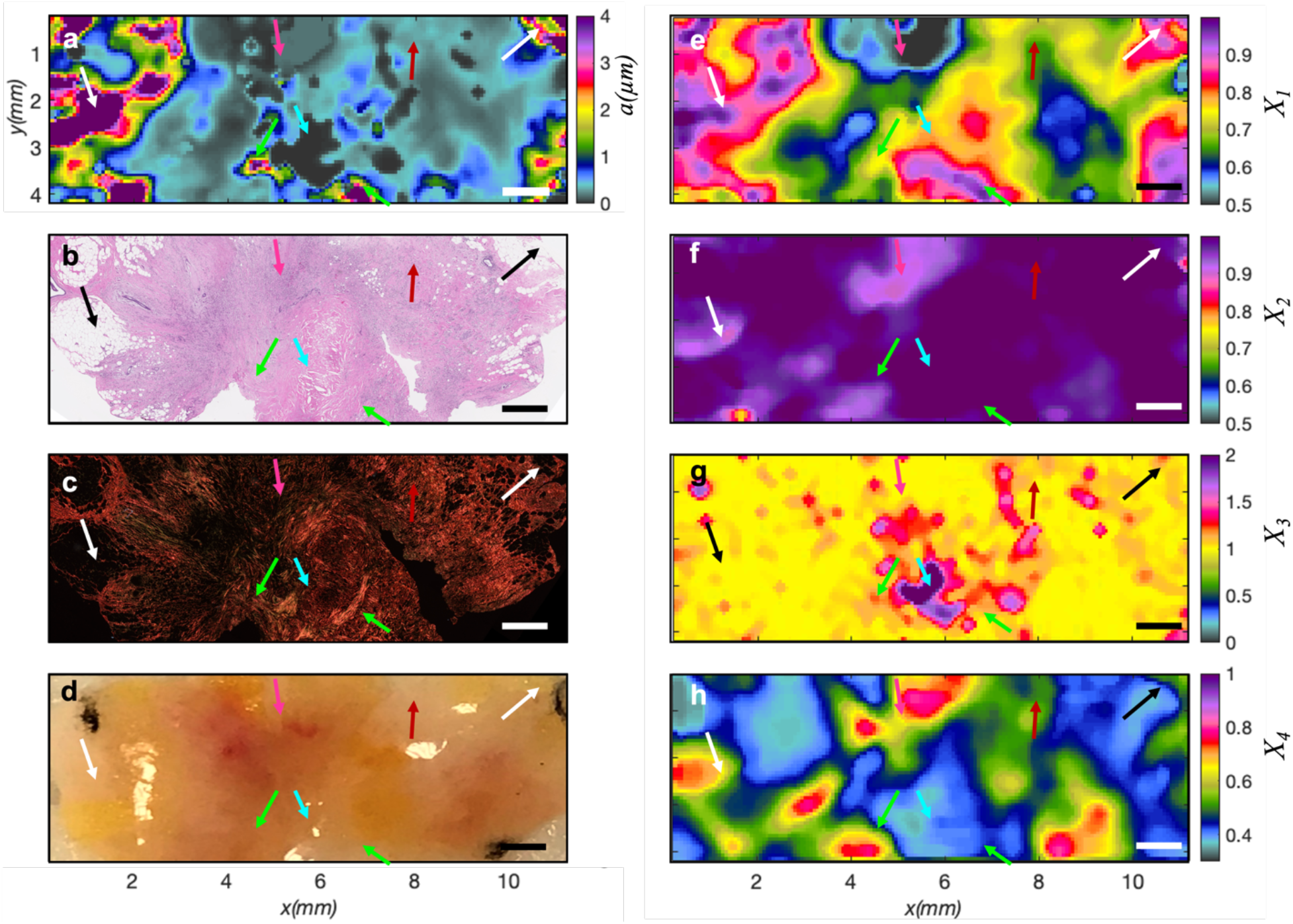
Particle size distribution in an invasive breast carcinoma microenvironment. **a** SPARSE map of *a*. Regions of increased particle size correspond to fat globules in the periphery (black & white arrows). The cyan to dark blue hues, corresponding to *a*:200nm-1μm, and denoted by maroon arrow, represent the regions rich in stromal fibers. The size drops to lower in the 100-200 nm range, when tumor cells are abundant (gray hues, magenta arrow). Islands of significant size drop to below 100 nm is highlighted by cyan arrow. These areas are surrounded by bands of *a*> 4μm, denoted by green arrows. **b** Matching H&E section entails the infiltrating tumor cells interspersed with stromal fibers taking up most of the tissue fragment, featuring a necrotic, scarred tissue in the middle, coincident with cyan arrows. In the peripheral areas, the tumor is invading the fibro adipose structures (black arrows). The tumor to stroma ratio is variable within different regions of H&E, with the maroon arrows corresponding to rich collagen content and magenta arrow pointing to cell-populated area. **c** PSR section reveals think, bundled collagen fibers, in bright orange and red hues, as denoted by green arrows in striking contrast with thin denatured green elastic debris broken down by tumor cells (cyan arrows). It also depicts the variations in tumor to stroma ratio across the tissue fragment. **d** Photograph displays the gross pathology of a human breast carcinoma tissue specimens. Presence of tumor vasculature is noted by the red hues (magenta arrow). **e** Spatial map of *X*_*1*_ displaying reduction of this metric within invasive areas that are populated by tumor cells and denatured collagen and are devoid of adipose and thick fibers. **f** Spatial map of *X*_*2*_ exhibiting lower values in cellular areas. **g** Spatial map of *X*_*3*_ is nearly 1 except for hollow areas in the PSR section (cyan arrow). **h** Spatial map of *X*_*4*_ showing reduction of this metric in the fibrous regions. Scale bars are 1 mm.

Histology of normal fibroadipose breast tissue typically involves a network of wavy collagen fibrils with radii ranging from 25 to 250 nm, interspersed with large adipocytes of approximately 20μm radius^37,38^. Figure 6 presents the sparse size map, the corresponding Hematoxylin & Eosin (H&E) and picrosirius-red (PSR) stained section, and the gross pathology of benign fibro-adipose breast tissue. The H&E slide reveals the arrangement of fibrous and adipose compartments, whereas the PSR highlights the presence of thin, undulating collagen fibrils within a single section (maroon arrows). SPARSE uncovers smaller structures below 1 μm in fibrous areas (maroon arrows), in striking contrast with larger structures exceeding 20 μm, coinciding with adipose regions (black or white arrows). Distinguished regions identified within the spatial maps of *X*_*1*_*-X*_*4*_ correspond to various tissue compartments. Notably, areas with diminished X_1_ align with fibrous tissue in both the H&E and PSR sections, likely due to smaller size of collagen fibrils in these areas. Conversely, elevated X_2_ coincides with fibrous areas in the gross pathology photo. In contrast, *X*_*3*_ remains nearly constant across the tissue, except for sporadic elevated specks that occasionally align with fibrous structures, and likely match the thinnest fibrillar structures (cyan arrows). Lastly, regions of reduced *X*_*4*_ correspond to the fibrous compartment, because smaller sizes are frequently associated with smaller *X*_*4*_. The individual *X*_*1*_*-X*_*4*_, capture only isolated features of the histology, and thus are not expected to present a one-on-one correlation with tissue ultrastructure.

However, it is intriguing that the size map (Fig. 6a) arising through identifying the local cluster and combining these metrics via the corresponding particle size estimation, yields an exquisite map of *a* variations that closely mirrors the histopathological features. In particular, the gray to blue hue regions in the size map, highlight submicron structures that correlate closely with the whereabouts of the collagenous component. On the other hand, the dark purple saturated areas in the size map, coincide with the adipose regions, suggesting that *a* ≥ 20μm in these areas. It is to be noted, however, that due to the finite beam spot size, scanning pitch, and the volume-integrated nature of the SPARSE measurement, in contrast to the single 7um thick histopathological sections, small microstructural differences are observed.

Next, SPARSE was further evaluated on an invasive ductal carcinoma specimen obtained from a patient with breast cancer (Fig. 7). The color bar range is reduced to 0-4μm to better visualize the smaller size scales emerging in the carcinoma compared to normal specimen. Presence of these significantly smaller sizes is likely due to the abundance of tumor cells and their sub-cellular structures, as well as the breakdown of collagen meshwork to ultrathin fibrils. The SPARSE map of Fig. 7a features a saturated region of larger particle size in the peripheral adipose region (*a*>4μm) as shown by the black & white arrows. Also featured in the map are cyan to navy blue regions, of *a*= 200nm-1μm, highlighted by maroon arrows that coincide with intra-tumoral stroma, and match the pink collagen hues in the H&E section and brighter red hues in the PSR image. SPARSE further reveals a region embedded in the tumor core that exhibits an abrupt size drop to below 100 nm, denoted by the cyan arrow. Comparison with the H&E-stained slide reveals the presence of the necrotic area devoid of cells that over time had transformed into scar tissue comprised of fissured collagen network. This region of significant reduced size is encircled by a few isolated islands of larger scatterers up to and exceeding 4μm. Examining the corresponding area within the PSR section reveals bundled thick collagen fibers surrounding the necrosed region, as highlighted by green arrows. Another conspicuous compartment, identified in the SPARSE map, features 100 nm<*a*<200 nm, as highlighted by the magenta arrow. The matching H&E section reveals the intermediate-grade tumor cells in this region, which invade in a trabecular pattern into the peripheral adipose tissue. Therefore, one would expect that larger size of tumor cell nuclei in this region, 1.5-2 times of an RBC, in the order of 6 μm, to dominate the size in this region^39,40^. However, close examination of corresponding H&E image and the gross photo suggest the presence of the tumor epithelium, intra-tumoral stromal fibers, and vasculature in this region. As such, speckle is formed not only by the large nuclei of tumor cells and the RBCs but also the cell membranes and organelles as well as denatured collagen fibrils. Therefore, the significant heterogeneity and polydispersity of the breast tissue, causes the SPARSE map to reflect the average particle size of the neighborhood. As in Fig 6 above, the spatial maps of *X*_*1*_, *X*_*2*_, *X*_*3*_, and *X*_*4*_ overall show variations with the tissue histopathology. For instance, *X*_*1*_ is higher in fibrous and adipose regions compared to tumor epithelium. This is particularly accentuated in the region pointed to by the magenta arrow. Reviewing all 4 metrics in this region suggests that the relative values of *X*_*1*_*-X*_*4*_ here (*X*_*1*_∼0.6, low; *X*_*2*_=0.85, low; *X3*∼1; *X*_*4*_=0.85 high) somewhat resembles the trends seen in the RBCs of whole blood above, in agreement with the presence of blood in gross pathology. However, the smaller scattering particles muddle the *X*_*1*_*-X*_*4*_, causing them to deviate from what is expected from larger RBCs or tumor cell nuclei. Conversely, in the stromal components, identified through the PSR image, *X*_*1*_ lies in 0.7-0.95 range, *X*_*2*_∼1, X_3_ varies between 1 and 2, and X_4_=0.5. This reveals significant heterogeneity in the particle size, consistent with the thick collagen fiber bundles interspersed with fissured and decomposed fibrils within the tumor stroma. In particular, specks of *X*_*3*_ *∼2* coincide with foci of collagen breakdown, whereas regions of maximized *X*_*1*_ match the thicker bundled collagen strands. Therefore, as seen in Fig. 6, *X*_*1*_*-X*_*4*_, while each capturing certain isolated aspects of tissue ultrastructure, represent only a coarse correlation with tissue morphology. Intriguingly, however, when these metrics are used to identify the local cluster and are substituted in the corresponding particle size estimation equation, the spatial distribution of *a* emerges that varies in an exquisite harmony with the histopathological landscape of the tumor tissue specimen.

## Discussion

Particle size characterization over a wide range of scales is a universal unmet need across various scientific disciplines, including food sciences, pharmaceutical industry, and clinical medicine. Conventional techniques like DLS and LD are typically limited to low-viscosity, dilute samples, offering minimal utility in biomaterials due to the complexity involved in detecting high-resolution angular scattering distributions or capturing light dynamics over extended time frames^4,12^. To ameliorate these deficits of traditional techniques, alternative approaches have been recently investigated. For instance, a low-cost miniaturized device was developed based on a spatial array of apertures with assorted diameters, each admitting distinct sets of scattering angles^41^. A machine learning model corrected for multiple scattering and predicted the size^41^. However, demonstrations were limited to measuring the median diameter of mono-dispersed large glass beads in the 13-125 μm range, while also requiring prior knowledge of bead concentration, and as a result, not conducive for biological specimens with smaller size scales and unknown particle concentrations or optical properties^41^. More recently, other groups demonstrated that the spatial autocorrelation of speckle informed the particle size distribution of powdered solid drugs in the 100-500μm range^13^. Nevertheless, this feature of laser speckle was most informative for sizing powdered solids of larger grain sizes spread on a surface, during processes such as drying, blending, and milling and presented limited applicability to biomaterials^13^. We and others had previously observed the feasibility to qualitatively estimate the size, through comparing the azimuth angle variations of *Î*_∥_ with an LUT^21,42^. However, this only allowed for semi-quantitative, discrete size estimation within the 250nm-2.5μm range, primarily distinguishing sizes slightly smaller or larger than the wavelength based on whether the scattering follows Rayleigh or Mie formalisms, and was limited to purely scattering samples with generally known refractive indices^21,42^.

Here we presented, for the first time, the SPARSE technique, that leverages spatial and time-dependent attributes derived from laser speckle patterns emerging due to interactions of polarized coherent light and the endogenous light scattering particles to characterize the size of particularities in a variety of tissues and biomaterials within the 10nm-10 μm range. This was achieved by developing a synthetic library of particle size and optical properties and the corresponding quantifiable metrics measured from the spatio-temporal attributes of polarized laser speckle patterns. Cluster analysis for the metrics identified 5 mutually exclusive regions, in each of which a tailored particle size estimation equation was devised to calculate the size as a function of the metrics. This particle sizing algorithm was then used to infer the size from the experimentally measured metrics of the polarized speckle.

The original synthetic library that supports the SPARSE prediction model, was developed assuming mono-dispersed spherical particles, exhibiting a *n*_*rel*_ =1.1 relative to the microenvironment. Therefore, SPARSE measurements in standard polystyrene microspheres given identical *n*_*rel*_ values, exhibited strong, statistically significant correlation with DLS. Deviations at *a<100 nm* occurred likely because the particle size estimation equation at this range, is primarily driven by *X*_*3*_. As explained earlier, all the metrics, including *X*_*3*_, depend on *a, μ*_*a*_ and *μ*_*s*_*’*. Yet, at smaller sizes, *μ*_*s*_*’* itself drastically varies by *a*. More specifically, for a given particle concentration, as *a* drop below 100 nm, the scattering cross section and in turn *μ*_*s*_*’* reduce significantly. Therefore, a competing effect emerges between the reduction of *a* and *μ*_*s*_*’* on *X*_*3*_, causing this metric to saturate at smaller sizes. Consequently, a large *X*_*3*_ may be equivocally interpreted as a moderate concentration of minute particles or a dense concentration of slightly larger scatterers, causing the estimated size to deviate from DLS measurements.

SPARSE was successful in characterizing the average size of spheroidal fat globules and protein micelles in milk samples. Retrospective review of refractive index variations in milk revealed that the fat globules indeed elicited a *n*_*rel*_*=1*.*1*^29^. However, protein micelles were more refractive^30^, and in principle could exhibit *X*_*1*_*-X*_*4*_ metrics that varied differently with size, compared to the synthetic library developed based on *n*_*rel*_*=1*.*1*, in turn hampering the utility of particle size estimation equation in milk. However, for *a*<500 nm, *X*_*1*_*-X*_*4*_ metrics and in turn the size estimation equations do not significantly change with *n*_*rel*_ variations (Supplementary Materials), suggesting that variations in *n*_*rel*_ does not compromise the applicability of the prediction model at the size ranges present in milk. The homogenization process likely reduced the variability of fat globules size distribution and in turn the polydispersity of milk samples, causing the SPARSE prediction model to remain applicable in milk phantoms. In addition, the ability of SPARSE to detect the few nm shifts in size caused by increasing the fat concentration from 0% to 4%, demonstrated the acute sensitivity of this approach in the 100-200 nm range, as validated through comparison with DLS measurements. On the other hand, DLS declared an average size that significantly correlated with SPARSE but also revealed the distinct sizes of protein and fat particle populations. Nevertheless, to conduct DLS measurements, samples were extensively diluted in deionized water, whereas SPARSE measured the size in untampered milk phantoms, suggesting the utility of this approach for monitoring the texture of food products composed of spheroidal particles of unknown concentrations.

Compared to polystyrene microspheres and milk, sizing RBCs in whole blood posed additional challenges, due to changes in both real and imaginary parts of *n*_*rel*_, and in turn *μ*_*a*_ and *μ*_*s*_*’*, concurrent to *a*. The challenge was readily apparent in isotonic blood, where the elliptical envelope of *Î*_∥_ resulted in *X*_*1*_=0.65, which in isolation would suggest a small *a*. However, by collectively considering this attribute in the context of other metrics (*X*_*2*_-*X*_*4*_), SPARSE successfully yields *a*=2.2 μm, which is close to the expected value of 2.8 μm^36^. It is to be noted that the lower *n*_*rel*_*=*1.03 of blood caused the scattering events to be increasingly forwardly directed and the optical paths of backscattered rays to be significantly longer compared to the above samples. This in turn enhanced the impact of absorption in terminating the longer optical paths, to an extent that was not recapitulated in the synthetic data based on MCRT assuming *n*_*rel*_*=*1.1. As saline concentration increases to 1.25% and 1.42%, absorption is reduced. Reduced absorption shifted the path length distribution towards longer paths. In addition, the lower *n*_*rel*_ made the scattering increasingly forwardly directed, which required more scattering events for the light to back-scatter and remerge at the surface. This resulted in significantly longer optical paths, which inherently emerged further away from the illumination center, causing the *Î*_⊥_ to expand radially, and *X*_*4*_ to enhance to nearly 1.4. As a result, erroneous presumption of the larger *n*_*rel*_=1.1 when predicting the size would correspond to *a* in the order of only a few 100s nm. Nevertheless, *n*_*rel*_ reduction was readily apparent from the envelope of *Î*_⊥_, given the shifted the maximum intensity from the envelope centroid to the middle of the leaflets. Therefore, we addressed the impact of *n*_*rel*_ variations by developing a synthetic library of *[X*_*1*_, *X*_*2*_, *X*_*3*_, *X*_*4*_], using a reduced *n*_*rel*_=1.03, and a whole new set of particle size estimation equations. Switching to this prediction model helped restore the SPARSE predictions at 1.25% and 1.42%, concentrations, where the reduced *n*_*rel*_ was readily inferred from the *Î*_⊥_. As solute concentration increased further, *n*_*rel*_ increased, causing the synthetic library and the particle size estimation equations, derived based on *n*_*rel*_=1.1, to be applicable to the experimentally measured metrics and trace the reduction in RBC size once again. While blood samples were significantly polydisperse, RBCs accounted for the majority of the speckle signal. However, RBCs are not spherical, and instead typically have a discoid shape. Yet, SPARSE was able to track the shift in the sphere-equivalent radii of RBCs in whole blood in its native state, likely because rich multiple scattering from numerous randomly oriented RBCs inherently averaged out the contribution of short and long dimensions, and shape features on the scattering signal. In contrast, DLS measurements were sensitive to shape. Our single angle, conventional DLS measurements conducted using parallel and perpendicular polarization filters, resulted in strikingly different size values, commensurate with the long and short dimensions of RBCs. Therefore, SPARSE measurements of blood were benchmarked against MADLS, and the measurements over multiple angles were averaged to permit comparison with SPARSE. Nevertheless, MADLS required diluting the blood with saline to a nearly transparent suspensions and took nearly 6 minutes to conclude for each concentration. We suspect that the overestimation of size in MADLS may have been a consequence of both the residual shape effects and RBC sedimentation during measurements.

As opposed to microspheres, milk, and blood, breast tissue specimens presented a more complex context given the variations in size, *n*_*rel*_, absorption, and particle concentration across the specimen. Nevertheless, tightly focusing the beam and scanning it across the sample minimized the influence of these variabilities within the illumination volume, causing the model to remain applicable at each scanning point. Furthermore, rich multiple scattering in tissue implied that the emergent speckle could be treated as the superposition of speckle patterns from different mono-disperse populations, causing the SPARSE prediction model developed using a synthetic library of mono-dispersed homogenous turbid medium, to remain resilient to polydispersity. This is evidenced by our results in normal fibroadipose tissue, which elucidate the characteristic size distribution of adipocytes and collagen fibrils, in their corresponding compartments identified in the H&E and PSR images. Compared to normal fibroadipose tissue, invasive ductal carcinoma presented a greater deal of size heterogeneity, as evidenced by histological analysis, which could likely complicate the size estimation.

Specifically, in such polydisperse turbid samples, photon path length distributions and in turn the metrics *X*_*1*_*-X*_*4*_ are modulated not only by the polarization status and average particle size, but also by polydispersity within the illuminated volume. Considering speckle patterns a simple superposition of signal due to subpopulations of small and large particles, *Î*_∥_ is expected to be influenced primarily by smaller particles which contribute shorter optical paths due to their isotropic scattering. As a result, *X*_*1*_ likely represents the smaller size populations, approximately within the *a*≤*λ* size range. This may be accentuated in biological tissue, where the scattering signal from numerous smaller particles overshadows the few larger scattering centers. Conversely, *Î*_⊥_ may be dominated by larger forwardly scattering particles in *a>λ* nm range, for which the longer optical paths cater to cross-polarized light. Thus, *X*_*2*_ and *X*_*4*_ could be biased towards larger particles in the population. Considering differences in the contributions of size scales to co- and cross-polarized speckle, *X*_*3*_ could present a different behavior in polydisperse and heterogeneous specimens. This is because the co-polarized speckle is contributed by shorter paths involving smaller particles. On the other hand, cross-polarized speckle is likely contributed by longer paths encountering larger particles. In addition, decay rates of g_2∥_(t), and g_2⊥_(t) at early and long times are modulated by faster and slower dynamics, elicited by smaller and larger particles, encountered in longer and shorter paths, respectively. Together, these suggest a time-variant log(g_2⊥_(t))/log(g_2∥_(t)), causing X_3_ to take on a wider range of values, beyond 1-1.8 observed in monodisperse samples, due to additional implications imparted by polydispersity.

Despite the additional implications of individual metrics in the context of polydisperse tissue, SPARSE size maps exhibited a detailed correspondence with histopathological features and varied in harmony with tissue compartments of varying morphologies. Fig. 7a demonstrates that the relative size differences nicely isolate histologically distinct regions in the highly heterogeneous breast carcinoma specimen. However, accurate absolute values were in some cases harder to estimate. For instance, the difficulty of distinguishing cell nuclei size is likely caused by the abundance of other organelles, membranes, and collagen fibrils. overshadowing the signal emanated from the nuclei within the illuminated volume. In the future, using a smaller beam spot and smaller scanning steps could likely increase the resolution and contrast of SPARSE measurements to permit isolating absolute values of different size populations.

The capabilities of SPARSE, demonstrated above, provide transformative opportunities across diverse disciplines. By enabling the assessment of the size distribution of nano-formulated drugs SPARSE could provide invaluable insight into drug stability and viability. In food science, SPARSE may be utilized to optimize milk processing to ensure enhanced product quality, texture, and nutritional value while maintaining food safety standards. This technology may be further advanced to accurately characterize the size variations of red blood cells in the context of various anemia and other hematological disorders. Through characterizing the size distribution of cells and their nuclei SPARSE could provide a surrogate biomarker for the more subjective and less quantitative tumor grade in cancer diagnosis. Taken together, SPARSE’s broad applicability across various fields facilitates size-based advancements, fostering innovation in industry and clinical medicine.

## Materials & Methods

### SPARSE Optical Setup

Figure 1 (a) displays the schematic diagram of the SPARSE optical setup. A polarized laser beam (Thorlabs, HNL210LB, 633 nm, 21 mW) is collimated and expanded to a beam diameter of 1 cm, passed through a focusing lens (Thorlabs, LA1986-A, Plano-Convex, AR coated, f= 124.6 mm), a beam splitter (Thorlabs, BS013, non-polarizing, beam splitter cube), resulting in a 10 μm spot of 5 mW on the sample surface. The backscattered light rays are collected and split by a second beam splitter in two different light paths and detected by two different high-speed CMOS cameras (Basler, ACa 2000-340 km, Germany). A pair of macro-lenses (Computar, MLH-10X) and linear polarizers focus the back-scattered light onto the CMOS sensor of the two camera. Adjusting the macro-lens f/# to 11 and magnification to 1X, results in a speckle size of S=2.44λ(1+M)f/#=34μm, and a one-dimensional pixel-to-speckle ratio of 5.7, ensuring sufficient spatial sampling, beyond Nyquist theorem^43^. The acquisition frame rate is selected according to the dynamics of the evaluated sample to ensure sufficient temporal sampling and adequate speckle contrast. The rotating mounts of the collection polarizers are adjusted so that the cameras acquire the co- and cross-polarized speckle patterns with respect to the illumination beam. To permit spatial mapping in heterogeneous biological tissues, the sample is placed in a holder and mounted on a 2-axis motorized stage (Newport, MFA-CC), controlled by a motion controller (Newport, ESP301). The sample is scanned in a serpentine pattern, with a scan pitch of 150 μm, to cover an area of a few mm^2^. At each scanned spot, speckle frames are acquired at 390 fps, for a sensor ROI of 1.2×1.2 mm, for 2 seconds. A custom-build C++ acquisition software enables adjusting the frame rate, exposure time, acquisition time, and frame size and synchronizes the motion controller and the cameras.

### Quantifying Size-Dependent Attributes of the Polarized Speckle Patterns

The entirety of the computation was carried out using MATLAB 2022a. Speckle frames were processed to obtain the diffuse reflectance profiles in the co- and cross-polarization states, *Î*_∥_(*x, y*) and *Î*_⊥_ (*x, y*), as follows:

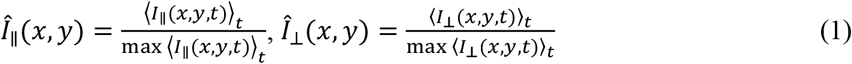

where, *I*_∥_(*x, y, t*) and *I*_⊥_ (*x, y, t*) refer to speckle intensities at location (x, y) and time instance t, and ⟨⟩_t_ denotes temporal averaging. In addition, speckle intensity autocorrelation curves for both co- and cross-polarizations are calculated according to:

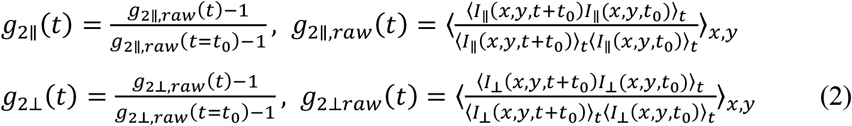

where, raw g_2_(t) curves are normalized to the speckle contrast^44^.

To calculate *X*_*1*_, *Î*_∥_(*x, y*) is contoured at 30% of its maximum and the inner and outer circles of the contour are identified. Radial averaging of *Î*_∥_(*x, y*) in the annular region between the two circles yields *Î*_∥_(*φ*), from which:

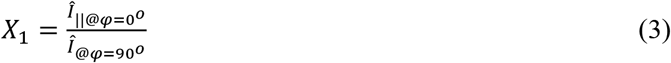

where *φ* stands for azimuth angle, and *φ* = 90 ° corresponds to the direction aligned with the polarization axis of the illumination beam. Likewise, *Î*_⊥_ (*x, y*) is contoured at 30% of its maximum, and the area and perimeter of the contour, A and P, are evaluated to calculate:

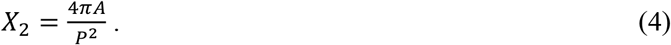

In addition, the differential decorrelation rate of co- and cross-polarized speckle frame series is quantified via:

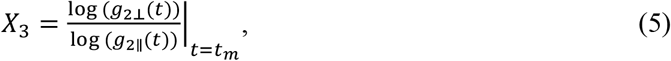

Where t_m_ is the instance in time where the temporal derivative of g_2_(t) curves are maximized. Finally, the average radius of the *Î*_⊥_ (*x, y*) contour is calculated as:

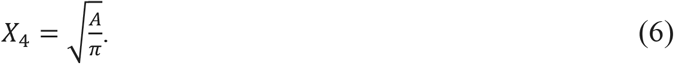

### SPARSE clustering analysis and particle size estimation

We exploited our previously developed polarization-sensitive correlation-transfer Monte-Carlo Ray tracing approach to simulate the propagation of a focused laser beam in a turbid media exhibiting *a* : 10nm -10 *μ*m, *μ*_*s*_*’*: 0.5-4 mm^-1^, and *μ*_*a*_: 0-70%μ_s_’ (Supplementary Methods)^19-21^. From the photon trajectories, the first- and second-order statistics, i. e. *Î*_∥_, *Î*_⊥_, g_2∥_(t), and g_2⊥_(t), of remitted speckle patterns and the corresponding size-dependent *[X*_*1*_, *X*_*2*_, *X*_*3*_, *X*_*4*_] metrics were simulated. K-means clustering was used to partition the synthetic library of *[X*_*1*_, *X*_*2*_, *X*_*3*_, *X*_*4*_] metrics (Supplementary materials) For each cluster, a step-wise multiple linear regression was used to derive the particle size estimation equation that yields *a* as a function of *[X*_*1*_, *X*_*2*_, *X*_*3*_, *X*_*4*_] metrics (Supplementary Methods). Pinpointing the of cluster centroids, and derivation of the particle size estimation equations is a one-time process. Once derived, these equations are readily applicable to predicting the scattering particle size from the experimentally evaluated *[X*_*1*_, *X*_*2*_, *X*_*3*_, *X*_*4*_] metrics. To this end, the *[X*_*1*_, *X*_*2*_, *X*_*3*_, *X*_*4*_] vector was assigned to a cluster with the centroid that exhibits the shortest Euclidean distance. Subsequently, the metrics were substituted in the particle size estimation equation, corresponding to the cluster, to yield the scattering particle size.

### Sample Preparation

Inert aqueous polarization microsphere suspensions of 10% weight fraction were obtained from the Bangs lab Inc. (NC, USA), exhibiting nominal radii in the range of 50 nm to 4800 nm (N=14). For microspheres smaller than 500 nm, 100 μm of the stock solution was mixed with 900 μm Glycerol, yielding a final polystyrene microsphere weight fraction of 1% in a 90% aqueous glycerol solution. Theoretical calculations, using Mie theory, suggest that *μ*_s_’ exhibits minimal variations around 1 mm^-1^, starting at 0.77 mm^-1^ for a=75 nm, reaching a peak of 1.08 mm^-1^ for a=100-200nm, and reducing to 0.81 mm^-1^ at a=400nm. For microspheres 500 nm and larger, the suspensions entailed 3% microsphere, 70% glycerol, and 27% water. The reduced viscosity of the solvent compensated for the increased particle size, and reduced the variability in the speckle fluctuations rate, permitting the same practical frame rate and acquisition duration across the size range. It also helped maintain biologically relevant *μ*_*s*_*’* values of 3 mm^-1^ for *a*=500 nm to 0.63 mm^-1^ for *a*=5μm. These *μ*_*s*_*’* values were only theoretical, and used only in the process of phantom design, but were not involved in the SPARSE particle size estimation. In practice, *μ*_*s*_*’* likely deviated from these baseline values due to experimental factors such as particle sedimentation or aggregation. The microsphere suspensions were placed in Eppendorf tubes and sonicated for 15 minutes to break down any potential aggregates. Subsequently, the samples were pipetted into custom-made 3D-printed cylindrical wells of 9 mm dia. and 1 cm depth. Speckle frame series were acquired at 1808 frames per second for 2 seconds in both parallel and perpendicular polarization states over 5 different points on the sample surface. Milk samples were obtained from a local grocery store (Wholefoods Market) at 0% (non-fat), 1% (low-fat), 2% (reduced-fat), and 4% (whole-milk) fat concentrations. The specimens were pipetted into the custom 3-D printed wells for SPARSE measurements.

One blue-top tube of whole blood specimen from a healthy donor was obtained from MGH Core hematology lab, using our IRB-approved protocol. A total of 11 samples were prepared by aliquoting 50 μl of whole blood Eppendorf tubes and spiking them with 10μl of saline with concentrations of 0.9-20%, corresponding to final saline concentrations of 0.9%-4.08% in blood. The samples were then pipetted into a custom 3D-printed chamber, with a clear polycarbonate imaging window. In an isotonic environment, i.e. a medium of the same saline molarity (0.9% concentration), normal human red blood cell (RBC) are expected to be discoid, with an indented center. Conversely, in a hypertonic solution, osmosis pressure expels the water from inside of the cell to the extracellular space, causing RBCs to shrink and exhibit a notched, crenated appearance^32^.

Freshly excised de-identified and discarded breast carcinoma and benign normal tissue from matching sites were obtained from the MGH surgical pathology suite following mastectomy procedures (MGH IRB#2011P000301). The specimens were stored in phosphate buffer saline at 4 °C and measured fresh within 24 hr of collection. The specimens were placed on top of wet gauze in a 35 mm petri dish and marked with black ink at four corners to facilitate co-registration with histology. The specimens were warmed up in a water bath to 37°. The petri dish containing the specimens was placed in a sample holder, connected to a 2-axis motorized stage for SPARSE measurement.

### Histological Analysis

Following SPARSE imaging, breast tissue specimens were placed in 10% formalin for at least 72 hours and treated with dissect-aid. The specimens were then embedded in paraffin, sectioned at 100 μm increments in depth (7 μm thick sections), and stained with hematoxylin and eosin (H&E), and Picrosirius Red (PSR), following routine protocols and Wellman photo pathology core. The H&E slides were digitized using Nano-zoomer whole slide scanner (2.0 HT, Hamamatsu, Japan). PSR slides were manually digitized using a circularly-polarized light microscope (BX43, Nikon, USA, 4x). Fiducial ink marks in the sample photographs and brightfield images were used to co-register the SPARSE maps with histology.

### Dynamic Light Scattering (DLS)

The particle size values of polystyrene microsphere suspensions, milk, and blood were verified through comparison with DLS measurements (Zeta Sizer Ultra, Malvern Instruments). For polystyrene microspheres, 5μl of the stock suspension was mixed with 1 ml in a spectroscopic cuvette. In case of milk samples, 100μl of specimens was mixed with 900 μl of deionized water in the cuvette. As for blood samples, 200 μl saline solutions of 0.9-20% concentrations were placed in 11 spectroscopic cuvettes and topped with an additional 1 ml of 0.9% saline, leading to total saline concentrations of 0.9-4.08%. A 5μl droplet of blood was added to each cuvette and mixed with the saline. The spectroscopic cuvettes were sealed and loaded in a DLS instrument (ZetaSizer Ultra, Malvern Instrument). DLS measurements were conducted at 173° back-scattering angle. Due to the discoid shape of RBCs, DLS measurements obtained under parallel and perpendicular polarizations differed significantly, corresponding to the short and long axes of the cells, highlighting their non-spherical nature. To account for the different RBC dimensions, the Multi-Angle DLS (MADLS) procedure was also conducted, allowing for averaging across dimensions and making the benchmark measurement more compatible with the shape-integrating nature of SPARSE.

## Acknowledgments

This work is funded by the financial support from National Institutes of Health through the following grants R21EB028951 (ZH), R01HL142272 (SKN), R01HL119867 (SKN), and the Air Force Office of Scientific Research grant FA9550-20-1-0063 (SKN). The authors thank Dr. Jie Zhao and team at the Photopathology Core of the Wellman Center for Photomedicine for histological processing of the tissue specimens.

## Author contributions

Z.H. and S.K.N. conceptualized the technique and proposed the optical instrument. Z.H., Z.Z., and S.K.N. designed the experimental methodologies. Z.H developed the MCRT simulation algorithm and developed the synthetic library of metrics. Z.H. and Z.Z. contributed to the development of the sample preparation procedures. Z.H. and S.K.N. developed the particle size estimation approach. Z.H. and N.L. evaluated the polystyrene microsphere, milk, blood and breast tissue specimens and conducted the data analysis and visualization. Z.H. and S.K.N. wrote the manuscript with contributions from all other authors. All authors contributed to the data analysis and interpretation of the results.

## Conflict of Interests

Z.H. and S.K.N. are listed as inventors on a U.S. Patent Application pertaining to the SPARSE approach. The remaining authors declare that they have no conflict of interest.

## Data and materials availability

All data needed to evaluate the conclusions in the paper are present in the paper and/or the Supplementary Materials. Additional data related to this manuscript may be obtained from the authors upon reasonable request.

