## Supplementary material for "Laser Speckle Particle Sizer (SPARSE) Informs the Size Distribution of Tissue Granularities": LSA_SupplementaryMaterial_Submitted.pdf

#### Supplementary Methods 1: Generating synthetic library of polarized speckle attributes and the quantitative metrics

We expanded our previously developed polarization-sensitive correlation transfer Monte-Carlo Ray Tracing (PSCT-MCRT) approach, to simulate first- and second-order statistics of the polarized speckle, namely $\hat{I}_{||}$, $\hat{I}_{\perp}$, $g_{2||}(t)$, and $g_{2\perp}(t)$ in turbid media[1]. The simulated media exhibited the following size range and optical properties: $a$: 10nm -10 $\mu$m, $\mu_s'$: 0.5-4 mm$^{-1}$, and $\mu_a$: 0-70%$\mu_s'$. Refractive indices of the particles and the surrounding medium were 1.59 and 1.45 respectively. This resulted in a relative refractive index mismatch of $n_{rel}$=1.1. For each $a$, Mie theory calculations were used to adjust the particle concentration to attain the desired $\mu_s'$ value. For each combination of $[a, \mu_s'=1\ mm^{-1}, \mu_a/\mu_s']$, a total of 1 million photons were traced in the medium, as they scattered off from one particle and collided with the next until finally remerged back at the surface. A simple geometric scaling was used to generalize the results to the entire $\mu_s'$ range. At each scattering event, Mie theory was used to relate the parallel and perpendicular polarized components of the back-scattered light as a function of the incident light using a matrix with diagonal entries of the S1 and S2 parameters as follows[2]:

$$\begin{bmatrix} E_{||s} \\ E_{\perp s} \end{bmatrix} = \frac{e^{jk(R-z)}}{-jkR} \begin{bmatrix} S_2(\theta) & 0 \\ 0 & S_1(\theta) \end{bmatrix} \begin{bmatrix} E_{||i} \\ E_{\perp i} \end{bmatrix} \quad (1)$$

Here, $E_i$ accounts for the incident field and $E_s$ represents the scattered electric field, where

$$E_i = E_0 e^{-jkz+j\omega} \quad (2)$$

In which the $E_0$ is the magnitude, $k$ is the wavenumber, $z$ is the traveling direction of the incident beam and $\omega=2\pi/\lambda$ is the angular frequency. S1 and S2 depend on the size parameter $x$, or the ratio of the particle size to wavelength, the complex index of refraction of the particle, and the scattering polar angle, $\theta$, through a series of Riccati-Bessel functions, the spherical Bessel functions, and the spherical Henkel functions[2]. Accordingly, during each scattering event, the direction of the photon changed in azimuthal, $\phi$, and polar angles, $\theta$, based on joint probability distribution function of:

$$P(\theta,\varphi) = I_0 S_{11}(\theta) + S_{12}(\theta)[Q_0\cos(2\varphi) + U_0\sin(2\varphi)] \quad (3)$$

Here $[I_0, Q_0, U_0, V_0]$ stands for the Stokes vector of the incident photon. The Stokes vector of the photon is updated using the single scattering function, as follows[1,3-6]:

$$\begin{bmatrix} I_s \\ Q_s \\ U_s \\ V_s \end{bmatrix} = \frac{1}{k^2R^2} \begin{bmatrix} S_{11} & S_{12} & 0 & 0 \\ S_{12} & S_{22} & 0 & 0 \\ 0 & 0 & S_{33} & S_{34} \\ 0 & 0 & -S_{34} & S_{33} \end{bmatrix} \mathrm{R}(\varphi) \begin{bmatrix} I_i \\ Q_i \\ U_i \\ V_i \end{bmatrix} \tag{4}$$

where

$$S_{11} = \frac{1}{2}(|S_2|^2 + |S_1|^2)$$

$$S_{12} = \frac{1}{2}(|S_2|^2 - |S_1|^2)$$

$$S_{33} = \frac{1}{2}(S_2^* S_1 + S_2 S_1^*)$$

$$S_{34} = \frac{1}{2} i(S_1 S_2^* - S_2 S_1^*) \tag{5}$$

The distance traveled between the successive scattering events is randomly distributed with an average value of $1/\mu_t$, where $\mu_t = \mu_s+\mu_a$, $\mu_s$ is the scattering coefficient and $\mu_a$ is the absorption coefficient. From these simulations, $\hat{I}_{||}$, $\hat{I}_{\perp}$, $g_{2||}(t)$, and $g_{2\perp}(t)$ are extracted for the said $[a, \mu_s'=1\ mm^{-1}, \mu_a/\mu_s']$, as [1,3-6]:

$$\hat{I}_{||} = \mathrm{IR} + \mathrm{QR} \tag{6}$$

$$\hat{I}_{\perp} = \mathrm{IR} - \mathrm{QR} \tag{7}$$

$$\mathrm{g}_{2||}^{\mathrm{MCRT}}(\mathrm{t}) - 1 = \left(\sum_{\mathrm{i}=1}^{\mathrm{n}} \mathrm{W_i}(\mathrm{I_i} + \mathrm{Q_i}) e^{-\frac{1}{3}\mathrm{k}^2\langle \Delta \mathrm{r}^2(\mathrm{t})\rangle \mathrm{Y_i}}\right)^2 \tag{8}$$

$$g_{2\perp}^{MCRT}(t) - 1 = \left(\sum_{i=1}^{n} W_i(I_i - Q_i) e^{-\frac{1}{3}k^2\langle \Delta r^2(t)\rangle Y_i}\right)^2 \tag{9}$$

Here, IR and QR are the total values of the first two elements of the stokes vectors on the surface, obtained by spatial binning of the Stokes vectors of the returning photons. In addition, $I_i$ and $Q_i$ are the first two elements of the Stokes vectors of the individual returning photons. Moreover, $W_i$ and $Y_i$ are the energy and momentum transfer of the individual returning photons.

The PSCT-MCRT algorithm was executed for a matrix of particle size values $a$, ranging from 10 nm to 10 µm, $\mu_s'$=1mm$^{-1}$, and $\mu_a$ ranging between 0 and 0.7 mm$^{-1}$, in 0.1 mm$^{-1}$ increments. A simple geometric scaling yields the photon trajectories for all possible combinations with $\mu_s'$=0.5-4mm$^{-1}$ range.

From this exhaustive set of simulated $\hat{I}_{||}$, $\hat{I}_{\perp}$, $g_{2||}(t)$, and $g_{2\perp}(t)$, we calculated the size-dependent metrics $[X_1, X_2, X_3, X_4]$ for each $[a, \mu_s', \mu_a/\mu_s']$ combination, as depicted in Fig. 1 of the manuscript. We further repeated these simulations by changing the $n_{rel}$ to 1.03 and 1.07, while keeping other variables the same, as depicted in Fig. S1 below.

**Supplementary Methods 2: Accommodating for refractive index variations**

To accommodate biological samples of lower refractive index variations such as blood, the PSCT-MCRT simulations was repeated for the same range of $a$, $\mu_s'$, and $\mu_a$ and reducing the $n_{rel}$ to 1.07

and 1.03. Supplementary Figures S1 and S1 display the simulated $\hat{I}_{||}$, $\hat{I}_{\perp}$ for a range of particle sizes, $a$, assuming $\mu_s'$=1*,* and $\mu_a$=0, and $n_{rel}$ values of 1.1, 1.07, and 1.03, respectively.

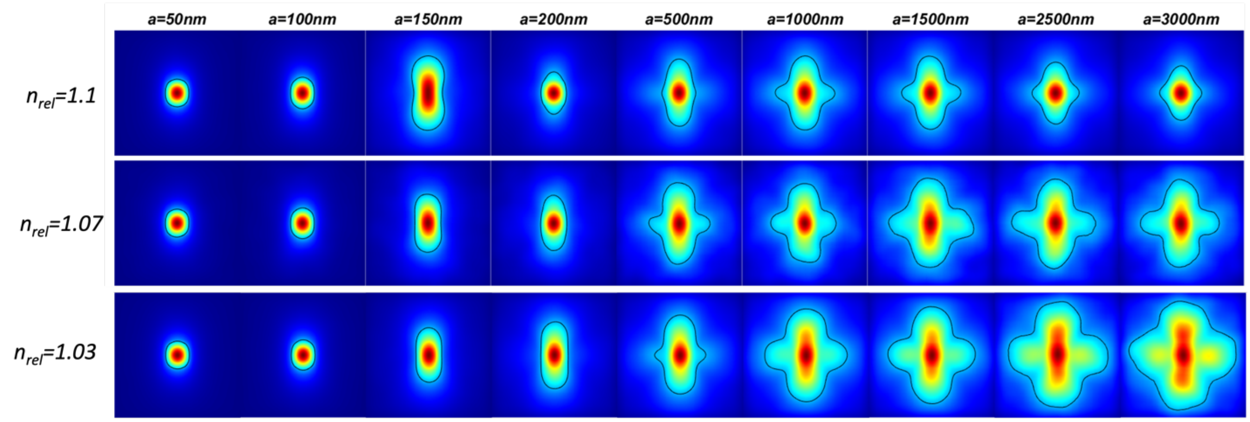


**Supplementary Figure S1. PSCT-MCRT simulation of $\hat{I}_{||}$, for various relative refractive index mismatch $n_{rel}$.** $\hat{I}_{||}$ envelopes are similar for different $n_{rel}$ values as long as a<500 nm. For larger particles, a lower $n_{rel}$ leads to an expanded $\hat{I}_{||}$ with relatively less prominent secondary pair of lobes perpendicular to the polarization axis.

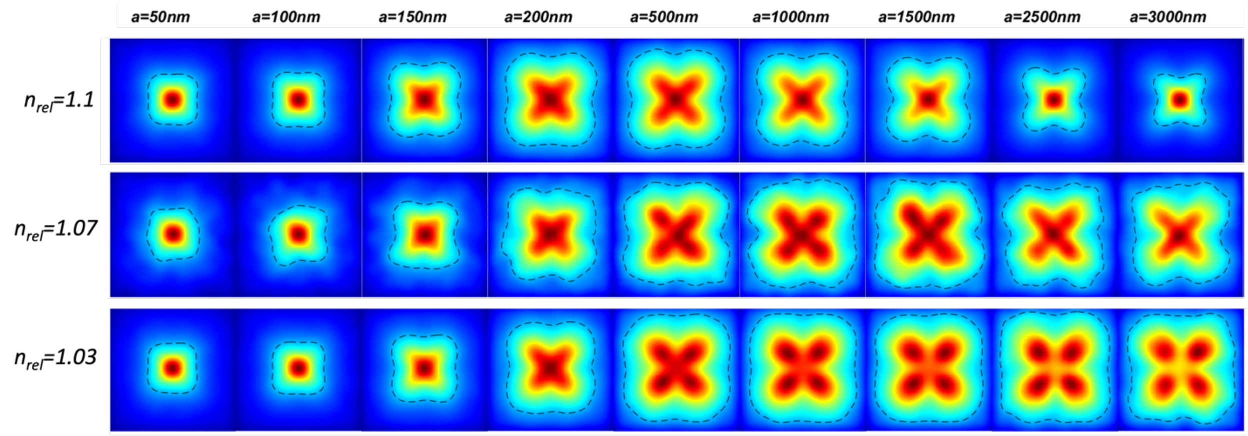


**Supplementary Figure S2. PSCT-MCRT simulation of $\hat{I}_{\perp}$, for various relative refractive index mismatch $n_{rel}$.** $\hat{I}_{\perp}$ envelopes are similar for different $n_{rel}$ values as long as a<500 nm. For larger particles, a lower $n_{rel}$, leads to an expanded $\hat{I}_{\perp}$. In addition, while the envelope remails clover-like, its circularity is reduced. Moreover, the intensity maximum shifts from the centroid to the middle of the leaflets. This trend will eventually stop for $n_{rel}$=1.07 at a>2500nm but continues for $n_{rel}$=1.03.

These additional sets of PSCT-MCRT simulations, and the resulting synthetic libraries of *[$X_1$, $X_2$, $X_3$, $X_4$]* metrics were used to develop two additional cluster analysis and particle size estimation equations. Supplementary Figure S3 displays the volume plots of $X_1$, $X_2$, $X_3$, and $X_4$ variations by the particle size, $a$, reduced scattering coefficient $\mu_s'$, and the normalized absorption coefficient, $\mu_a/\mu_s'$, as well as their cross sections at $\mu_s' = 1\ mm^{-1}$, assuming $n_{rel}$=1.1, 1.07, and 1.03. In addition, Figure S4 displays the cluster assignment and the scatter diagram of $a$ vs $X_1$, $X_2$, $X_3$, $X_4$, color-coded with the cluster number. Note that depending on $n_{rel}$, clusters correspond to different segments of $\mu_s'$, $\mu_a/\mu_s'$, and $a$ space. Also, please note that, one may not fully isolate *a,* $n_{rel}$*,* and $\mu_s'$. This is because $n_{rel}$ and $a$ determine the range of possible values for $\mu_s'$. For instance, if $n_{rel}$ =1.03, $\mu_s'$ may not exceed 2 $mm^{-1}$ since for such a low refractive index mismatch, the sample may not exhibit a very strong scattering even at higher scattering particle concentrations. Therefore, the cluster assignments displayed in supplementary Fig. S4 for $n_{rel}$=1.03 and $n_{rel}$=1.07 are displayed for $\mu_s'$,=0.5-4 mm$^{-1}$ only to remain consistent with the initial particle size estimation equations. The synthetic library of *[X1, X2, X3, X4]* metrics was used to develop the cluster analysis and the particle size estimation equation of SPARSE, as detailed later.

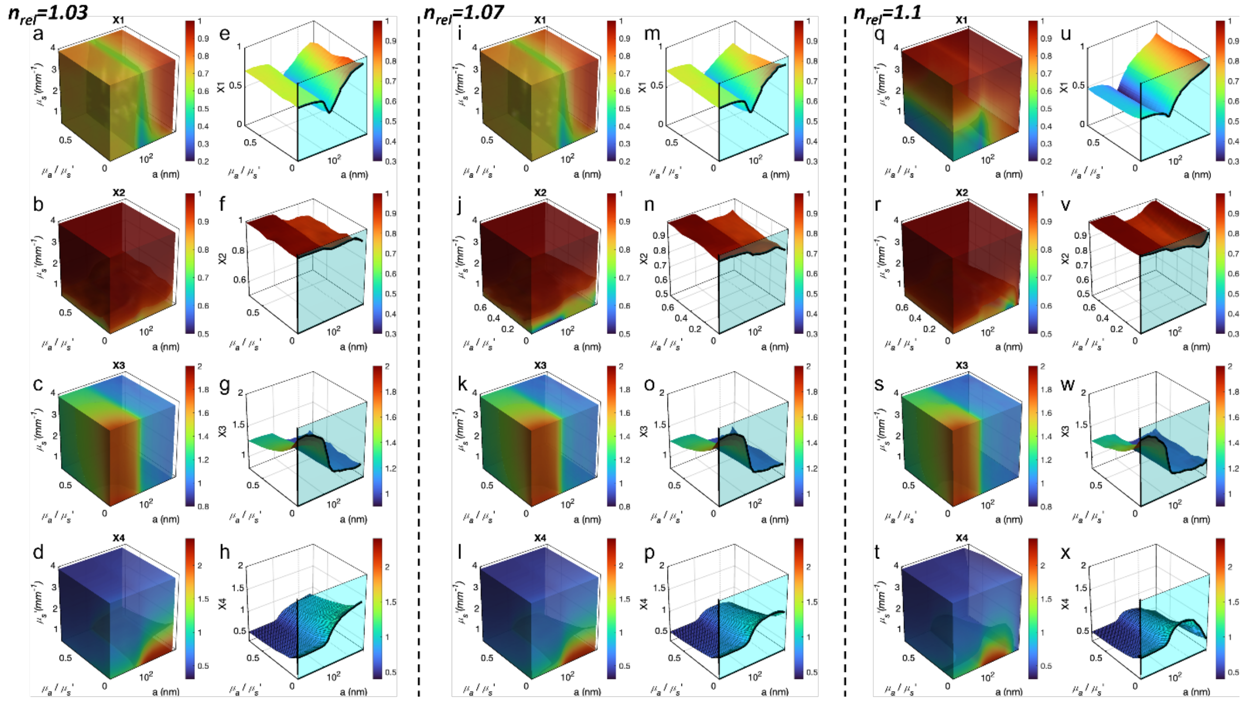


**Supplementary Figure S3. Synthetic library of size-dependent attributes, created by a Polarized CT-MCRT.** a-d Volume plots of $X_1$, $X_2$, $X_3$, $X_4$ variations by the particle size, a, reduced scattering coefficient $\mu_s$', and the normalized absorption coefficient, $\mu_a$ /$\mu_s$', obtained assuming **$n_{rel}$=1.03**. e-h Cross-section of the volume plots displaying $X_1$, $X_2$, $X_3$, $X_4$ for a typical $\mu_s$'=1 mm$^{-1}$, displaying the variations of the metrics with the particle size, and absorption, obtained assuming $n_{rel}$=1.03. The wall plot, highlighted by a cyan background corresponds to $\mu_s$'=1 mm$^{-1}$ and $\mu_a$=0. i-l Volume plots of $X_1$, $X_2$, $X_3$, $X_4$ variations by the particle size, a, reduced scattering coefficient $\mu_s$', and the normalized absorption coefficient, $\mu_a$ /$\mu_s$', obtained assuming **$n_{rel}$=1.07**. m-p Cross-section of the volume plots displaying $X_1$, $X_2$, $X_3$, $X_4$ for a typical $\mu_s$'=1 mm$^{-1}$, displaying the variations of the metrics with the particle size, and absorption, obtained assuming $n_{rel}$=1.07. The wall plot, highlighted by a cyan background corresponds to $\mu_s$'=1 mm$^{-1}$ and $\mu_a$=0. q-t Volume plots of $X_1$, $X_2$, $X_3$, $X_4$ variations by the particle size, a, reduced scattering coefficient $\mu_s$', and the normalized absorption coefficient, $\mu_a$ /$\mu_s$', obtained assuming $n_{rel}$=1.1. u-x Cross-section of the volume plots displaying $X_1$, $X_2$, $X_3$, $X_4$ for a typical $\mu_s$'=1 mm$^{-1}$, displaying the variations of the metrics with the particle size, and absorption, obtained assuming **$n_{rel}$=1.1**. The wall plot, highlighted by a cyan background corresponds to $\mu_s$'=1 mm$^{-1}$ and $\mu_a$=0.

### Supplementary Methods 3: Investigating the impact of blood absorption on the metrics.

Supplementary Fig. S5 displays the PSCT-MCRT simulated $\hat{I}_{||}$, $\hat{I}_{\perp}$, for varying $\mu_a/\ \mu_s'$, assuming the RBC size $a$=2.8 $\mu m$, its relative refractive index compared to plasma $n_{rel}$=1.03, and the expected reduced scattering coefficient $\mu_s'$ =1 mm$^{-1}$. These simulations reveal that increased absorption prunes the longer paths, pushes the $\hat{I}_{||}$ to the double-lobed pattern, and shrinks the $\hat{I}_{||}$, $\hat{I}_{\perp}$. Consequently, both $X_1$ and $X_4$ are reduced in response to absorption. The simulated $\hat{I}_{||}$, $\hat{I}_{\perp}$ at $\mu_a/\ \mu_s'$=0.3 closely resemble the experimentally evaluated $\hat{I}_{||}$, $\hat{I}_{\perp}$ of isotonic whole blood in Fig. 5 of the manuscript.

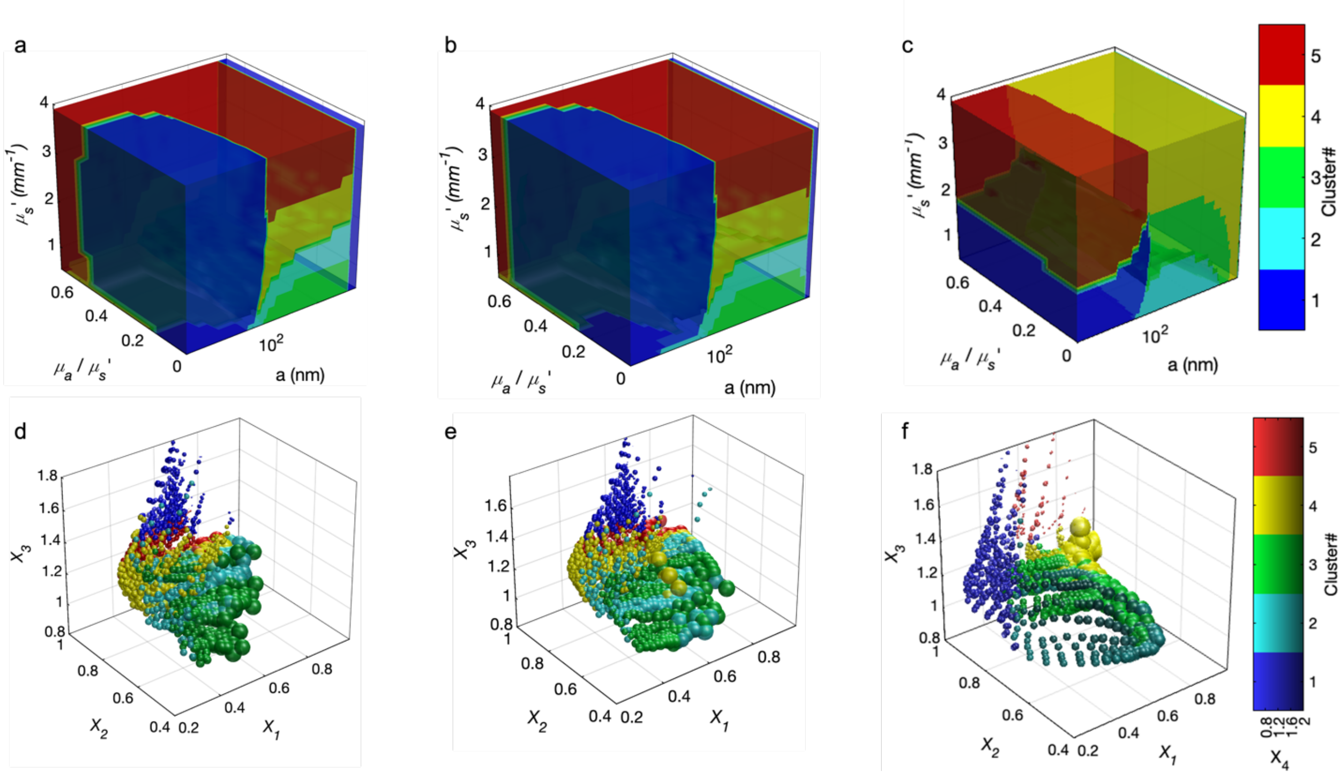


**Supplementary Figure S4. Cluster analysis of speckle metrics and scatter plots of particle size vs metrics for various $n_{rel}$. a-c** Cluster analysis of [$X_1$,$X_2$,$X_3$, $X_4$] metrics, obtained assuming $n_{rel}$ =1.03, 1.07, and 1.1, respectively. The clusters partition the [$a$, $\mu_a$, $\mu_s'$] space to 5 distinct clusters. **d-f** Scatter plot of the cluster assignments in the [$X_1$,$X_2$,$X_3$, $X_4$] space, obtained assuming $n_{rel}$ =1.03, 1.07, and 1.1, respectively. The 3 axes of the plot correspond to $X_1$, $X_2$, and $X_3$, while $X_4$ is depicted by varying the luminescence of the cluster color. The sizes of spherical markers are proportionate to the particle volume, i.e. $\sqrt[3]{a}$. A tailored particle size estimation equation that formulates $a$ as a function of $X_1$, $X_2$, $X_3$, $X_4$ is obtained for each cluster, and each $n_{rel}$.

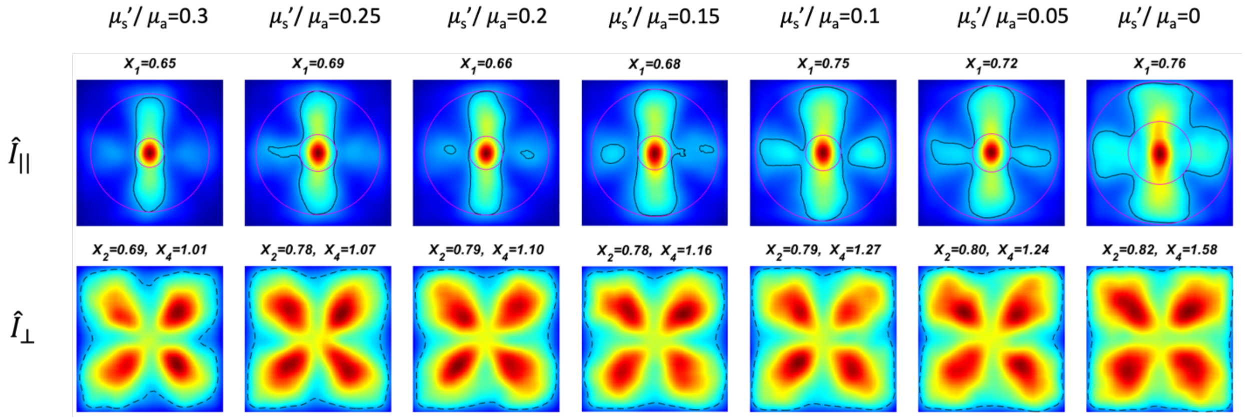


**Supplementary Figure S5. PSCT-MCRT simulation of $\hat{I}_{||}$ and $\hat{I}_{\perp}$ assuming an $n_{rel}$=1.3, and with varying $\mu_a$/ $\mu_s'$ ratios**. Increased absorption reduces the prominence of secondary pair of lobes perpendicular to the polarization axis in $\hat{I}_{||}$ , casuing it to transform from a four-leaved pattern to the double-lobed form. In addition, increased absorption causes both $\hat{I}_{||}$ and $\hat{I}_{\perp}$ to shrink, radially.

**Supplementary Methods 4: Developing the K-means clustering analysis & particle size estimation equations**

When attempting to develop particle size estimation equation, we noted that the small and large particles exhibit different dependencies on $[X_1, X_2, X_3, X_4]$. For instance, while in turbid media of smaller size scales and weak scattering, *a* exhibited an inverse linear relation with $X_1$, at larger size scales and rich scattering, the relationship was direct and exponential. The variability of the relationships between metrics $[X_1, X_2, X_3, X_4]$ metrics and *a* across the size scales, and additional dependence on optical properties precluded a single equation to estimate *a* from these metrics across all the size ranges and for materials of different optical properties. In another words, it appeared that multiple tailored particle size estimation equations govern the relationship between $[X_1, X_2, X_3, X_4]$ metrics and *a,* depending of the size range and turbidity levels.

For this reason, we employed clustering techniques to identify mutually exclusive regions within the $[X_1, X_2, X_3, X_4]$ metrics and *a* space, enabling the development of tailored particle size estimation equations for each region. To determine the optimal number and boundaries of these regions, we performed K-means clustering, a widely used method that partitions data into a specified number of clusters by minimizing the variance within each cluster[7]. K-means is particularly useful for its simplicity and efficiency in handling large datasets, making it well-suited for our objective of identifying distinct regions within the multidimensional space[7]. To calculate the K-centroids, first the centroids are randomly selected or initialized. Subsequently, each $[X_1, X_2, X_3, X_4]$ point is assigned to the nearest centroid. The centroids of the clusters are then updated as the mean of all points in the cluster. This process is repeated iteratively until the centroids stabilize or the maximum number of iterations is reached. The performance of the clustering approach is evaluated using the silhouette score, which measures how well each data point fits within its cluster compared to others. If most points have a high silhouette value, then the clustering solution is appropriate. Our analysis suggested that the silhouette score, is optimized when the space is divided into five clusters. These clusters effectively partitioned the simulated $[X_1, X_2, X_3, X_4]$ space into five distinct regions, each corresponding to a different region in the $[a, \mu_s', \mu_a/\mu_s']$ space.

It is important to note that our cluster analysis is based solely on the synthetic $[X_1, X_2, X_3, X_4]$ vectors, independent of the corresponding $[a, \mu_s', \mu_a/\mu s']$ values used to generate these metrics. Figure S4 presents the color-coded clusters within the $[a, \mu_s', \mu_a/\mu s']$ and $[X_1, X_2, X_3, X_4]$ spaces for various $n_{rel}$ values of 1.03, 1.07, and 1.1.

A few interesting observations can be made by reviewing the regions correspond to the clusters in the $[a, \mu_s', \mu_a/\ \mu_s']$ space. For instance, assuming $n_{rel}$=1.1, as displayed in Fig. S4 c, clusters #1 and #5 both roughly corresponds to turbid media with a<100 nm and $\mu_a/\ \mu_s'$ changing in the full 0-0.7 range, with the distinction that in cluster #1 $\mu_s'$<2 $mm^{-1}$ and in cluster#5 $\mu_s'$>2 $mm^{-1}$. In other words, if cluster analysis of experimentally evaluated $[X_1, X_2, X_3, X_4]$ suggest that the sample belongs to cluster#1, one can readily infer that a<100 nm and that the sample is less turbid. On the other hand, clusters 2,3, and 4 encompass turbid media with generally larger particles, and varying levels of turbidity. It is to be reiterated that these observations are merely retrospective, and that the k-means clustering was performed on the synthetic library of $[X_1, X_2, X_3, X_4]$ metrics irrespective of simulation variables $[a, \mu_s', \mu_a/\ \mu_s']$ used to generate the polarized laser speckle attributes. Table S1 lists the cluster centroids for $n_{rel}=1.1$.

**Table S1.** Cluster centroids.

| Cluster# | $X_1$ | $X_2$ | $X_3$ | $X_4$ |
|---|---|---|---|---|
| 1 | 0.43 | 0.94 | 1.36 | 0.79 |
| 2 | 0.61 | 0.71 | 1.05 | 1.85 |
| 3 | 0.68 | 0.84 | 1.05 | 1.08 |
| 4 | 0.89 | 0.99 | 1.06 | 0.58 |
| 5 | 0.84 | 1.0 | 1.53 | 0.52 |

Subsequently, a step-wise regression analysis was used to obtain the tailored particle size estimation equations specific to each cluster, through adding or removing the metrics and their interaction terms, starting from a constant model. The iterative stepwise approach identified the main effect of each of [$X_1$, $X_2$, $X_3$, $X_4$] metrics and their interaction terms to yield the most accurate model and minimize the mean square error. When devising the equation, it turned that a linear regression best suited cluster#1, i.e. less turbid media of small particle size. However, the particle size estimation equation was of exponential form for the remaining 4 clusters. The coefficients of particle size estimation equations corresponding to individual clusters, and assuming $n_{rel}=1.1$, are listed below:

**Table S2.** The coefficient of particle size estimation equations correspond to clusters# 1-5. The coefficients corresponding to individual attributes and the interaction terms are listed.

| Cluster# | Intercept | $X_1$ | $X_2$ | $X_3$ | $X_4$ | $X_1{\times}X_2$ | $X_1{\times}X_3$ | $X_1{\times}X_4$ | $X_2{\times}X_3$ | $X_2{\times}X_4$ | $X_3{\times}X_4$ |
|---|---|---|---|---|---|---|---|---|---|---|---|
| 1 | 439.69 | 0 | 0 | -254.97 | 0 | 0 | 0 | 0 | 0 | 0 | 0 |
| 2 | 3.65 | 0.97 | -1.85 | 0 | -0.95 | 0 | 0 | 0.60 | 0 | 0.64 | 0 |
| 3 | 6.52 | 0.65 | -6.27 | 0 | -2.18 | 2.73 | 0 | -0.44 | 0 | 2.53 | 0 |
| 4 | -182.73 | 108.45 | 151.58 | 132.09 | 14.05 | -83.30 | -13.73 | -15.46 | -107.67 | 31.48 | -31.95 |
| 5 | 126.28 | -119.87 | -123.62 | -2.29 | -89.99 | 120.07 | 0 | 0 | 0 | 94.05 | 0 |

The above table indicates that the coefficients of the particle size estimation equation, corresponding to the contribution of each of the *[$X_1$, $X_2$, $X_3$, $X_4$]* metrics and their interaction are different in each cluster. In other words, for each cluster, the particle size is determined from a different sets of metrics and interactions terms . Therefore, while all 4 metrics are used to identifying the cluster #, they are different weighted in the particle size estimations equations of individual clusters, as listed below:

$$Cluster\#1 \quad \hat{a} = 439.69 - 254.97X_1 \tag{10}$$

$$Cluster\#2 \quad \hat{a} = 10^{3.65+0.97X_1-1.85X_2-0.95X_4+0.6X_1X_4+0.64X_2X_4} \tag{11}$$

$$Cluster\#3 \quad \hat{a} = 10^{6.52+0.65X_1-6.27X_2-2.18X_4+2.73X_1X_2-0.44X_1X_4+2.53X_2X_4} \tag{12}$$

$$Cluster\#4 \quad \hat{a} = 10^{\left\{\begin{matrix}-182.73+108.45X_1+151.58X_2+132.09X_3+14.05X_4-83.3X_1X_2- \\ 13.73X_1X_3-15.46X_1X_4-107.67X_2X_3+31.48X_2X_4-31.95X_3X_4\end{matrix}\right\}} \tag{13}$$

$$Cluster\#5 \quad \hat{a} = 10^{126.28-119.87X_1-123.62X_2-2.29X_3-89.99X_4+120.07X_1X_2+94.05X_2X_4} \tag{14}$$

### Supplementary Data:

**Table S3** Nominal value of polystyrene microsphere radii value, characterized by the manufacturer, together with our SPARSE and DLS size measurements (Average size: μ, standard deviation: σ), along with theoretically calculated $\mu_s'$ values.

| *a (nm) (nominal)* | $\mu_{DLS}$ *(nm)* | $\sigma_{DLS}$ *(nm)* | $\mu_{SPARSE}$ *(nm)* | $\sigma_{SPARSE}$ *(nm)* | $\mu_s'$ ($mm^{-1}$) |
|---|---|---|---|---|---|
| *72.5* | *72.96* | *3.94* | *51.47* | *2.74* | *0.77* |
| *97.5* | *95.95* | *5.87* | *93.95* | *1.21* | *1.08* |
| *149* | *143.33* | *9.21* | *92.27* | *28.24* | *0.96* |
| *199* | *187.58* | *18.7* | *236.69* | *5.43* | *1.07* |
| *251* | *229.02* | *35.0* | *414.42* | *9.25* | *0.97* |
| *299* | *236.56* | *41.54* | *321.56* | *4.42* | *0.92* |
| *408* | *359.97* | *33.85* | *315.71* | *8.17* | *0.81* |
| *502* | *519.58* | *45.04* | *1482.99* | *44.13* | *3.05* |
| *1010* | *1211.7* | *163.51* | *1159.53* | *8.11* | *2.06* |
| *1505* | *1545.4* | *115.60* | *1500.36* | *34.20* | *1.62* |
| *2090* | *2098.7* | *111.93* | *1712.23* | *32.39* | *1.3* |
| *2900* | *2778.7* | *116.85* | *2235.57* | *481.58* | *1.07* |
| *4800* | *4677.7* | *763.49* | *5575.44* | *1852.27* | *0.63* |

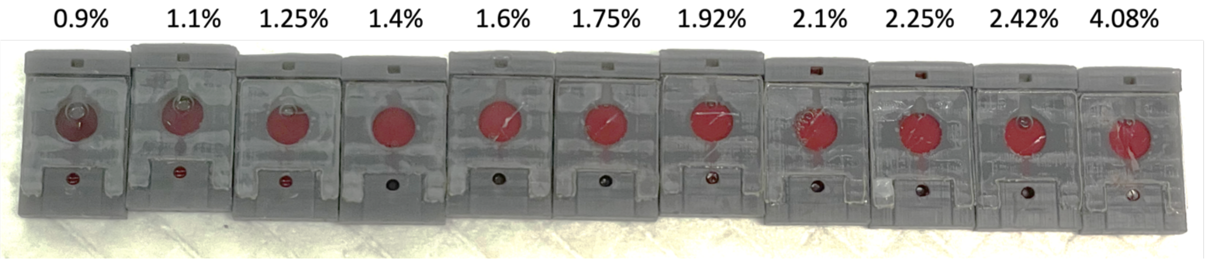


**Supplementary Figure S6. Whole blood specimen, spiked with saline solutions of different solute concentrations.** As the salt concentration increases, the red color becomes increasingly brighter and more vibrant, indicating reduced absorption and increased scattering.

### *References*

1 Hajjarian, Z. & Nadkarni, S. K. Estimation of particle size variations for laser speckle rheology of materials. *Opt Lett* **40**, 764-767 (2015). https://doi.org:10.1364/OL.40.000764

2 Hergert, W. & Wriedt, T. *The Mie Theory, Basics and Applications*. (Springer Berlin, Heidelberg, 2012).

3 Hajjarian, Z. & Nadkarni, S. K. Evaluation and correction for optical scattering variations in laser speckle rheology of biological fluids. *PloS one* **8**, e65014 (2013).

4 Hajjarian, Z. & Nadkarni, S. K. Correction of optical absorption and scattering variations in laser speckle rheology measurements. *Optics express* **22**, 6349-6361 (2014).
5 Ramella-Roman, J. C., Prahl, S. A. & Jacques, S. L. Three Monte Carlo programs of polarized light transport into scattering media: part I. *Optics express* **13** (2005).
6 Ramella-Roman, J. C., Prahl, S. A. & Jacques, S. L. Three Monte Carlo programs of polarized light transport into scattering media: part II. *Optics express* **13** (2005).
7 Kaufman, L. & Rousseeuw, P. J. *Finding groups in data : an introduction to cluster analysis*. (Wiley, 1990).
